# Unraveling Regulatory Feedback Mechanisms in Adult Neurogenesis Through Mathematical Modelling

**DOI:** 10.1101/2025.03.12.642763

**Authors:** Diana-Patricia Danciu, Filip Z. Klawe, Alexey Kazarnikov, Laura Femmer, Ekaterina Kostina, Ana Martin-Villalba, Anna Marciniak-Czochra

## Abstract

Adult neurogenesis is defined as the process by which new neurons are produced from neural stem cells in the adult brain. A comprehensive understanding of the mechanisms that regulate this process is essential for the development of effective interventions aimed at decelerating the decline of adult neurogenesis associated with ageing. Mathematical models provide a valuable tool for studying the dynamics of neural stem cells and their lineage, and have revealed alterations in these processes during the ageing process. The present study draws upon experimental data to explore how these processes are modulated by investigating regulatory feedback mechanisms among neural populations through the lens of nonlinear differential equations models. Our observations indicate that the time evolution of the neural lineage is predominantly regulated by neural stem cells, with more differentiated neural populations exerting a comparatively weaker influence. urthermore, we shed light on the manner in which different subpopulations govern these regulations and gain insights into the impact of specific perturbations on the system.

## 1. INTRODUCTION

Adult neurogenesis is the process by which mature neurons are generated from neural stem cells (NSCs) throughout adulthood. In mammals, adult neurogenesis takes place in two main regions of the brain: the dentate gyrus of the subgranular zone in the hippocampus and the ventricular-subventricular zone located along the walls of the lateral ventricles [1-3]. In this process, NSCs transition from a quiescent to an active state, enabling self-renewal and differentiation. This results in the generation of transient amplifying progenitors (TAPs), neuroblasts (NBs), and ultimately neurons. A schematic of these transitions among the neural populations is shown in Fig. 1A. With ageing, the number of mature neurons decreases, which may lead to impaired cognitive function [4-7]. This phenomenon has been reviewed in [8, 9], .Furthermore, it has been found that the number of neural stem cells declines with age in both the ventricular-subventricular zone [10-12] and the dentate gyrus of the adult hippocampus [13, 14], (for reviews, see,). As neural stem cells represent the pinnacle of the neural hierarchy and give rise to the entire neural lineage, interventions aimed at counteracting the effects of ageing could potentially have the greatest impact if they targeted stem cells. Consequently, the study of the dynamics of neural stem cell populations and their lineage is of fundamental importance for advancing our understanding of cognitive function in the context of ageing.

**Figure 1:**
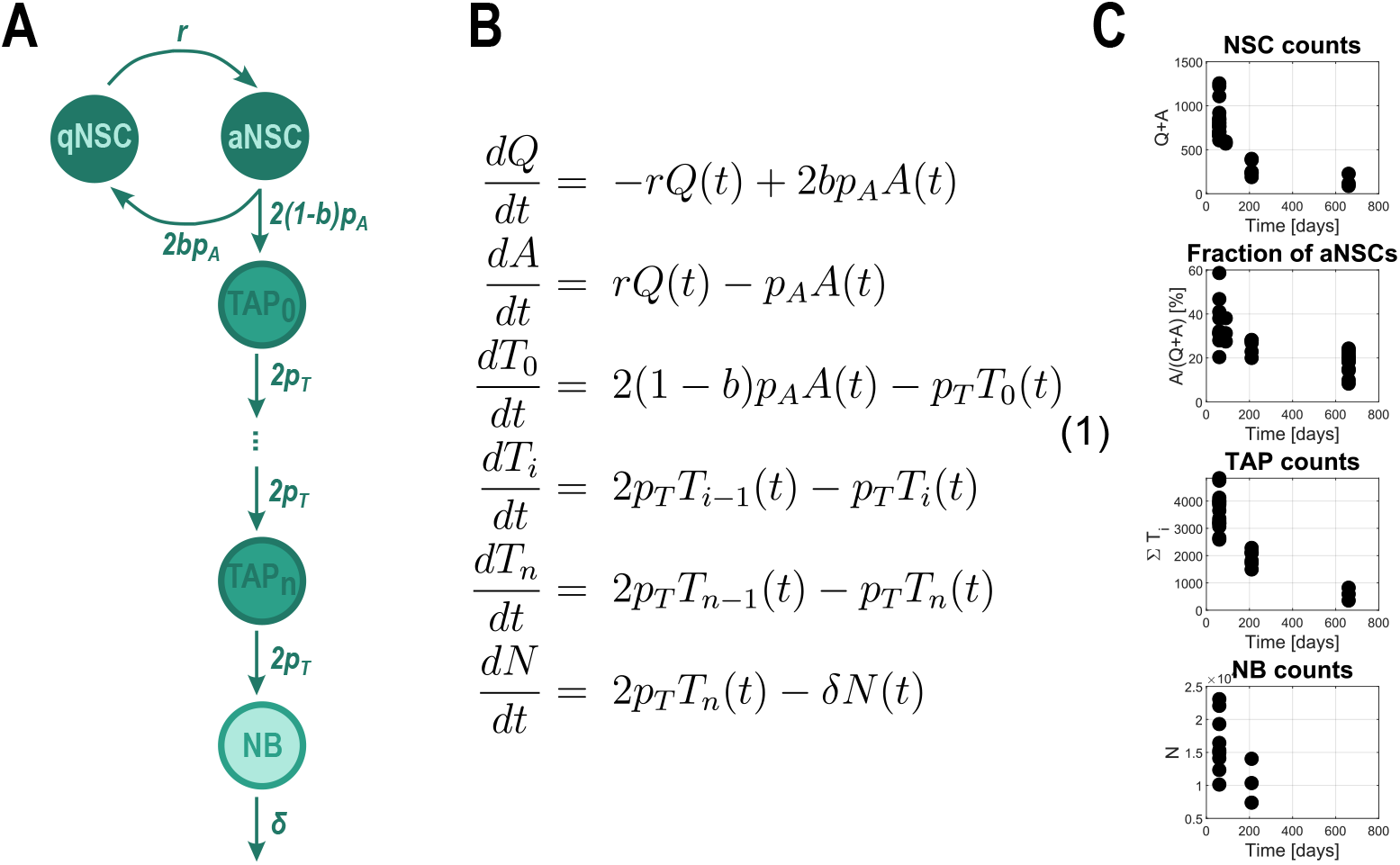
Mathematical model and data of cell population dynamics: **A**. Schematic of the transitions among neural cell types, represented in the mathematical model. The arrows and rates depicted represent the inflow from the previous compartment. **B**. Mathematical model. System of ODEs describing the time dynamics of the neural populations depicted in A. The number of amplification steps *n* = 3, corresponding to 4 TAP compartments *T*_*i*_. **C**. Existing data from [12, 36]: total number of NSCs; fraction of active cells among label retaining cells, confirmed to be a good approximation of the fraction of active neural stem cells among all neural stem cells; total number of TAPs and total number of NBs in the ventricular-subventricular zone. Each data point corresponds to one mouse.

As experimental studies have their limitations, mathematical methods can be employed to shed light onto processes that lead to the NSC dynamics described by the available data (Fig. 1C). Mechanistic mathematical models of the neural populations, involving non-autonomous linear ordinary differential equations (ODEs) with time-dependent system parameters, have facilitated uncovering trends in the time evolution of these parameters. Applied to the dentate gyrus of the adult hippocampus [14, 15] and to the ventricular-subventricular zone [12], they found that NSCs spend increasingly more time in quiescence as they progress with age. However, such models do not allow us to infer the regulatory mechanisms driving this behaviour, nor how different types of perturbations alter the time course of system parameters and, consequently, of cell populations. One of the key questions is how to realistically model system parameters in a way that enables a better understanding of the underlying regulatory processes.

These regulatory feedbacks are governed by cell-cell interactions through an elaborated network of signalling pathways. Different cells produce or interact with signalling molecules and can regulate activation, self-renewal, differentiation or proliferation. For example, it has been reported that, in zebrafish, non-glial neural progenitors similar to active NSCs and TAPs in mammals laterally inhibit the activation of quiescent NSCs by upregulating Notch ligands at the cell surface [16, 17]. Additionally, in mice it has been proposed that NSC quiescence is regulated by their immediate progeny through similar Notch paracrine signals [18], as well as diffusible ones such as the neurotransmitter ABA [19]. Notch has also been suggested to promote NSC proliferation while maintaining them in an undifferentiated state, in other words promoting self-renewal [20]. In contrast, the Ascl1 gene was found to be expressed at varying levels by proliferating lineage cells (aNSCs and TAPs) and to play a critical role in the activation of quiescent NSCs. Additionally, its inhibitor HES is expressed by quiescent cells and promotes the maintenance of quiescence. These are both tightly regulated by Notch expression, which plays an essential role in maintaining quiescent NSCs [21]. Further signals (among others) expressed in the neural lineage and involved in its regulation include bone morphogenetic protein (BMP), interferon (IFN) and Wnt. In particular, an interplay between canonical and non-canonical Wnt has been reported, with both playing a role in regulating system parameters.

A very large array of signals from the neural lineage is involved in coordinating the dynamics of NSCs and their neural lineage, with intricate interactions, whose study is extremely complex and outside the scope of this paper. These signals are further influenced by and interact with inputs from non-neural niche cells [22]. Ependymal, endothelial cells and astrocytes are some of the non-neural cell types from the neurogenic niche that contribute to regulating neurogenesis. Additionally, inflammatory signals such as Interferons (IFN) are not only involved from within the niche, but also from outside, for example through microglia or T-cells from the cerebrospinal fluid [23]. Overall, adult neurogenesis dynamics are regulated by a complex web of interactions among many different signalling pathways and cell types (Table 1), whose balance changes with ageing, and can be disrupted in neurological diseases [23-25]. Nevertheless, as a first step, before setting out to untangle this web, it seems advantageous to gain insights into where the driving combined signal originates and how it guides adult neurogenesis. This concept justifies the scope of our work on developing and employing nonlinear ODE models, with the aim of reducing the complexity and focusing on the basic driving actors in adult neurogenesis, their source and target.

**Table 1:** Overview of reported regulatory mechanisms.

| Function | Promoting | Inhibiting | Key signals | References |
| --- | --- | --- | --- | --- |
| Activation of qNSCs ($r$) | aNSCs, TAPs, Ependymal, Astrocytes, Microglia | qNSCs, aNSCs, TAPs, NBs, Astrocytes, Endothelial, Microglia | Wnt, EGF, FGF2, BMPs, Notch, Shh, TGF-β, GABA | [16, 20, 23, 25–29] |
| Self-renewal of aNSCs ($b$) | qNSCs, aNSCs, Astrocytes, Ependymal, Endothelial | TAPs, NBs, Astrocytes, Microglia | Notch, Wnt, EGF, FGF2, Shh, VEGF, BMPs | [20, 23, 25–28] |

**Table 2:** Summary of fixed and estimated parameters for each type of data.

| Experiment |  | Fixed | Estimated |
| --- | --- | --- | --- |
| WT | | $p_A, p_T, \delta, NSC_0$ | $r_0, K, b_0, \beta$ |
| TMZ | young | $r_0, K, b_0, \beta, p_A, \delta, NSC_0$ | $p_T, d, \rho$ |
| | old | $K, b_0, \beta, p_A, \delta, NSC_0$ | $r_0, p_T, d, \rho$ |
| IFNAGR KO | | $p_A, \delta, NSC_0$ | $r_0, r_1, K, b_0, \beta, p_T$ |

The present paper investigates regulatory feedbacks among cells of the neural lineage that lead to the observed dynamics. Using experimental data from wild-type (WT) mice, as well as complementary data from two perturbation settings, the mathematical model yields insights into the mechanisms governing the neural lineage. Our strategy is as follows: Based on previous models that identify temporal variability in key model parameters, we first determine the general form of the feedback function. We then identify this function using model-based data analysis, combining deterministic parameter estimation with Bayesian approach to uncertainty quantification. Finally, guided by the shape of the feedback function, we propose a specific molecular implementation that reproduces both wild-type data and data from perturbation experiments.

## 2. RESULTS

### 2.1. Neural lineage model and the motivation for feedback regulation

The model presented in this study builds upon previously established frameworks describing the population dynamics of different cell types [12, 14]. The main ingredients of the mechanistic model correspond to the transitions among the neural lineage compartments and, in particular, among the NSC subpopulations (depicted in Fig. 1A and defined by the ODE system (1) in Fig. 1). NSCs can exist in two exclusive states: active, which are actively cycling, and quiescent, which are not in the cell cycle. Quiescent NSCs (qNSCs) can become active at a rate *r* to enter the cell cycle and transition into the active compartment. An active NSC (aNSC) can divide at a rate *p*_*A*_ into either two qNSCs (symmetric self-renewal) or into two TAPs (symmetric differentiation) to advance further along the differentiation path. The balance between self-renewal and differentiation is modelled via the so-called fraction of self-renewal *b*. This represents how many progeny of NSCs are NSCs themselves, and at the single-cell level can be thought of as a probability that the NSC self-renews. Asymmetric divisions can also be encompassed in the self-renewal parameter without any change in the model equations (an explanation can be found in 30] and 31]). TAPs are considered to perform multiple amplification steps [32] at a rate *p*_*T*_ before finally dividing into NBs, which can exit the compartment at a rate *δ*, for example by death or by migrating to another brain region. According to biological evidence, a number of *n* = 3 amplification steps is assumed [33-35], corresponding to four TAP compartments *T*_*i*_, *i* ∈ {0, …, 3}. The equations (1) describing these assumptions are defined in Fig. 1B [12, 30].

The question is how to model the system’s parameters, and in particular, how to link them to signalling pathways that regulate cellular processes in order to understand systemic control. Our approach to this problem builds on insights gained from previous models demonstrating that models with constant parameters fail to reproduce the wild-type dynamics [12]. Instead, the activation rate of quiescent stem cells must vary over time to match experimental observations. Perturbation experiments further revealed a compensatory role of the time-varying self-renewal fraction during cell division, suggesting it helps buffer age-related changes in activation (Fig. 2A). These time-dependent parameters were inferred by fitting data under specific assumptions about the functional form of activation and self-renewal rates. Fig. 2A compares the dynamics of total stem cell numbers and the fraction of active cells, along with parameter trajectories, for scenarios in which either both parameters are constant (“no ageing”), one is time-dependent (“decreasing activation” or “increasing self-renewal”), or both vary over time (“regulating both”). In a related study, Dabelow et al. [37] applied optimal control theory to explore alternative trajectories of these parameters and concluded that a decreasing activation rate is the most influential driver of the observed ageing dynamics. However, these time-dependent models lack mechanistic insight into how such regulatory changes are implemented in the system. As a result, they fail to reproduce recovery dynamics following injury, which depend critically on the internal state of the system at the time of perturbation [12]. Furthermore, Harris et al. 15] showed that increasing the heterogeneity of the quiescent stem cell population—by modelling it as a mixture of shallow and deep quiescence states with distinct activation rates—cannot replace the need for time-dependent activation. This finding underscores that cellular heterogeneity alone is insufficient to explain the observed dynamics and highlights the necessity for system-level regulation mediated by signalling molecules.

**Figure 2:**
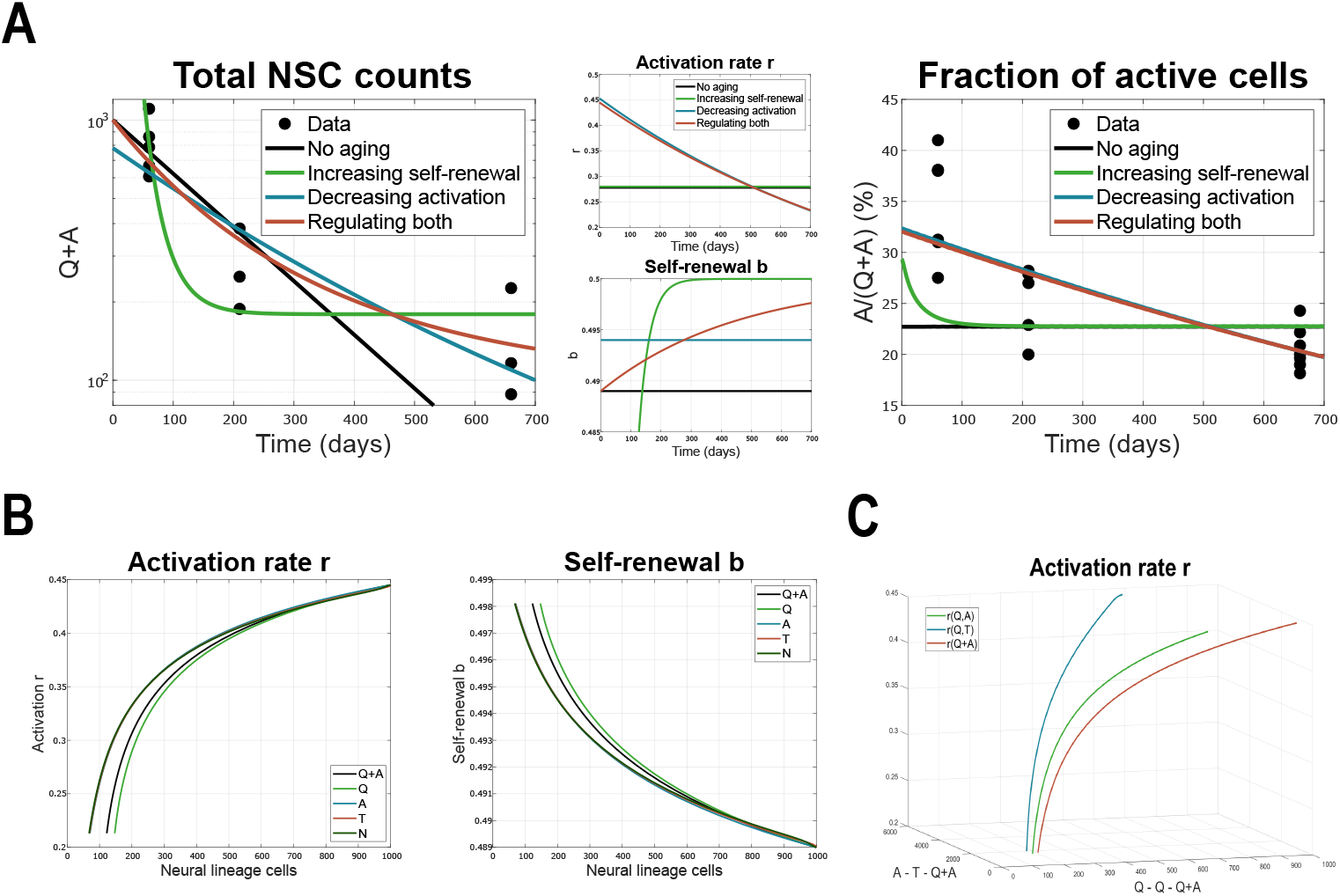
Identifying the time-dependence of system parameters **A**. Results of the various scenarios considered in [12] and reviewed in [30]. The system parameters are given by time-dependent functions 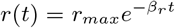 and 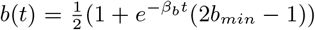 introduced in [12, 36], which are basic examples of decaying activation and increasing self-renewal. Plots adapted from [30]. **B**. System parameters, i.e. activation rate *r* and fraction of self-renewal *b* from the linear model plotted against values of various neural lineage populations (scaled up to 1000 cells). **C**. 3D plot of the activation rate *r* with respect to various combinations of neural cell subpopulations.

### 2.2. Identifying the shape of feedbacks

To uncover the regulatory feedbacks underlying the behaviour observed in experimental data, we employ nonlinear ODE models. The model structure is defined by the system of equations (1), as shown in Fig. 1B. In contrast to previous approaches, the system parameters—namely the activation rate *r* and the self-renewal fraction *b*-are now modelled as functions of neural lineage subpopulation sizes. It is important to note that what is often termed “self-renewal” in biological literature corresponds, in our framework, to the combined effect of *r* and *b*, two processes that are challenging to disentangle experimentally. For clarity, we refer to the parameter *b* as “self-renewal” throughout this study, while the combined effect of *r* and *b* is termed “effective self-renewal,” following the terminology introduced in [30].

How exactly do activation and self-renewal depend on the cellular context? By plotting parameter estimates from the linear models against the corresponding NSC counts over time, we observe that the activation rate *r* increases with the number of NSCs, whereas the self-renewal fraction *b* decreases. Notably, experimental data show that all neural lineage subpopulations decline over time (Fig.1C), suggesting that both *r* and *b* respond similarly to different subpopulation sizes. This is further illustrated in Fig.2B, which shows comparable parameter dynamics relative to various neural cell populations.

It is convenient to assume an activating Hill-like dependence of the activation rate *r*, and an inhibitory Hill-like dependence of the self-renewal fraction *b*, on cell population size. In other words, the system parameters can be described by Eq. (2),

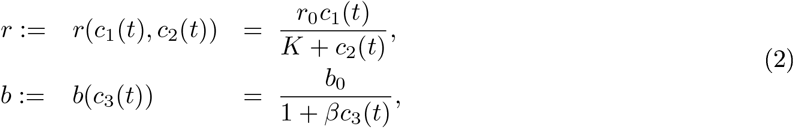

where *c*_*i*_ represent some cell subpopulation of the neural lineage, the parameters *r*_0_ and *b*_0_ govern the amplitude, and *K* and *β* tune the shape of the Hill functions. The same Hill-like behavior of *r* is observed whether *c*_1_ and *c*_2_ correspond to the same neural subpopulation that promotes activation, or to different neural subpopulations that regulate activation in opposing manners. This similarity can be observed in Fig. 2B and 2C, respectively.

In the following, we present model identification based on a range of experimental settings, including wild-type neurogenesis and perturbation experiments. These insights guided the search for a specific signalling pathway consistent with the identified regulatory structure.

### 2.3. Model selection for wild-type adult neurogenesis uncovers quiescent NSCs as the dominant promoter of activation

We consider various scenarios corresponding to different subpopulations inserted in the *c*_*i*_ of our formulas (2). After performing weighted-least-squares parameter estimation (described in Methods) to find how each scenario compares with the experimental data from WT mice, we examine the results of scenarios grouped by the regulator of self-renewal. Each panel in Fig. 3 depicts the comparison among data and different hypotheses for feebacks in the activation rate of qNSCs, for self-renewal regulated by individual subpopulations (qNSCs, aNSCs, total TAPs, NBs or total NSCs).

**Figure 3:**
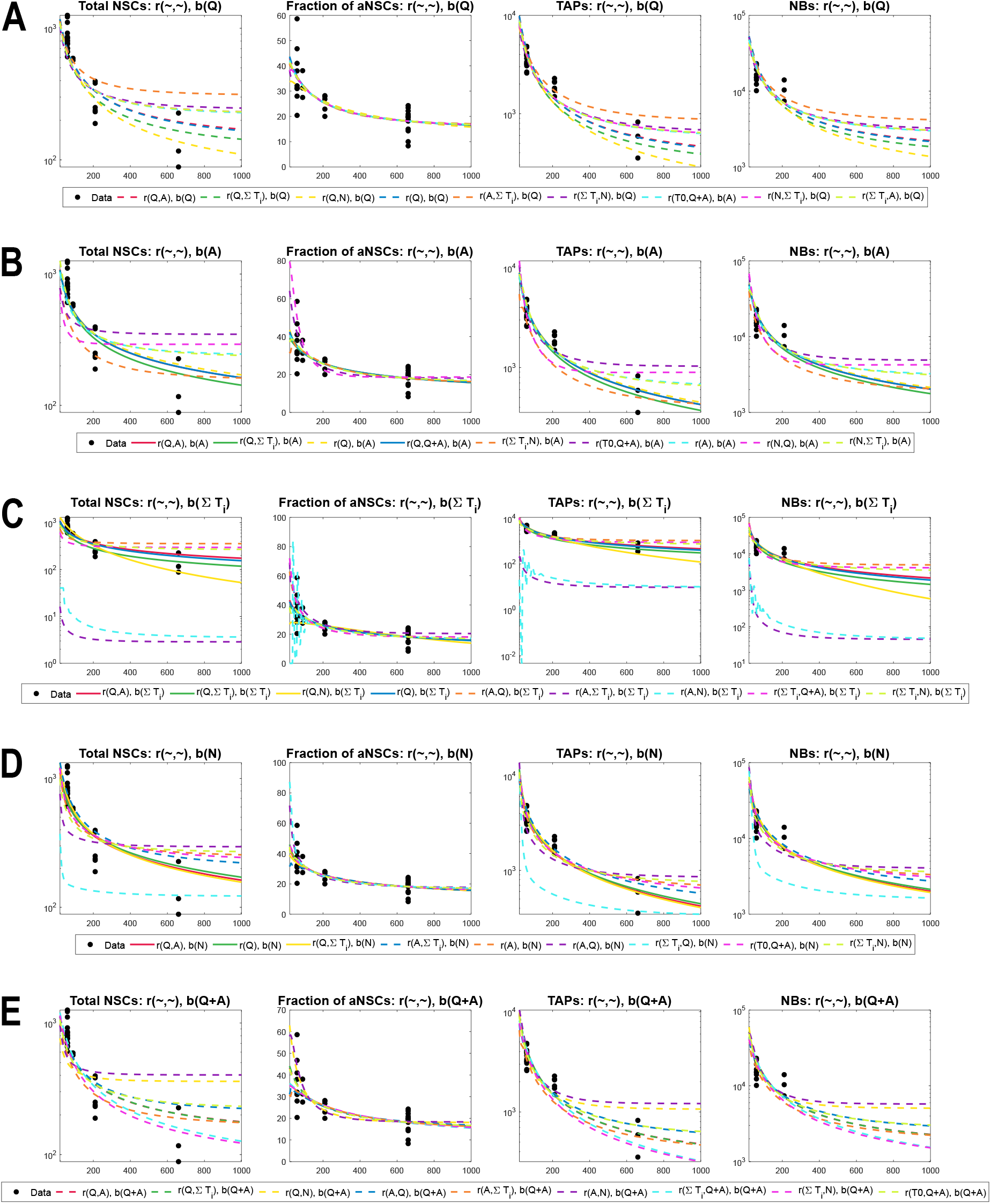
Comparison among various scenarios for the regulation of activation rate, grouped by the inhibitor of the self-renewal: **A**. Self-renewal inhibited by qNSC, *b*(*Q*). **B**. Self-renewal inhibited by aNSC, *b*(*A*). **C**. Self-renewal inhibited by T s, *b*(Σ*T*_*i*_). **D**. Self-renewal inhibited by N s, *b*(*N*). **E**. Self-renewal inhibited by total number of NSCs, *b*(*Q* + *A*). s a notation rule, *r*(*c*_1_, *c*_2_) corresponds to activating ill functions (2) in which *c*_1_ promotes (is in the numerator) and *c*_2_ inhibits (denominator) *r*. ere NSC counts represent *Q*(*t*) + *A*(*t*), fraction of active corresponds to *A*(*t*)*/*(*Q*(*t*) + *A*(*t*)), TAP counts is considered the sum 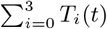 and NB counts is given by *N* (*t*). Solid lines represent scenarios in which the estimated parameter *b*_0_ *<* 0.5 and the system only admits the trivial equilibrium. Dotted lines correspond to scenarios in which *b*_0_ *>* 0.5 and thus a stable positive steady state exists. X-axis represents Time, i.e. the age of mice in days. Y-axis is on log-scale in all except the plot depicting the fraction of active cells. For readability, in the plot legends we write the multivariate functions *r*(*c*_1_) := *r*(*c*_1_, *c*_1_).

To compare models, we compute the Akaike Information Criterion corrected for small sample sizes (AICc). This approach assigns each model a score that reflects its ability to explain the data while penalising complexity to avoid overfitting. Differences in these scores allow for the identification of a “best-scoring” model, while the corresponding Akaike weights indicate the relative likelihood that a given model is truly the best among those considered (see Methods for details). owever, due to the very limited number of experimental data points, the AICc scores are subject to considerable uncertainty and may shift substantially with additional data. Therefore, rather than selecting a single model, we focus on the top-scoring candidates, testing their predictions through perturbation experiments and using their structure to explore biologically plausible mechanisms involving signalling molecules.

A first observation from Fig. 3 is that the subpopulation that inhibits activation is located further along the differentiation path than the one promoting the activation. Interestingly, the scenarios in which qNSCs promote activation capture the dynamics of the experimental data the best. Some hypotheses that align with the summary in Table 1 also provide an acceptable fit, but not as good as the former. Furthermore, the mathematical models clearly show that the quiescent NSCs do not have an inhibitory effect on both the activation rate and the fraction of self-renewal (Supplementary Figure 1A). These modelling results suggest that, even though evidence of qNSCs maintaining quiescence and proliferating cells promoting activation exists (Table 1), this is not the primary mechanism involved. Instead, the balance between the combined effect of neural subpopulations and thus signalling pathways tips towards a dominant effect from qNSCs in driving activation.

There exist a large number of possible combinations of the cell types *c*_*i*_ from Eq. (2), some more plausible than others. Previous experimental studies have proposed that activation of qNSCs is promoted by aNSCs and TAPs with the aim of sustaining neurogenesis, and inhibited by NBs. In contrast, it is also reasonable to consider that the activation rate *r* may be inhibited by proliferating cells, such that the quiescent population is conserved when enough cells are already cycling, as was reported in zebrafish [16]. Intuitively, an activation rate that increases with the number of qNSCs is also a reasonable assumption: an abundance of qNSCs allows them to activate, as the main goal of the system is to produce neurons, not only conserve its stem cell pool. An alternative may be that all NSCs, quiescent or active, express the same driving signals for activation or self-renewal, so that the system does not distinguish the source of the feedback. As far as the regulation of self-renewal is concerned, other biological systems such as hematopoiesis [38, 39] suggest that it is inhibited by cells at the end of the differentiation spectrum (mature cells, analogous to our NBs). Such strategy seems logical if the purpose is to replenish a small NB population, since it is more efficient to first produce more stem cells that then proliferate at the same rate as before, than to proliferate faster and differentiate more [38, 4]. Self-renewal may also be inhibited by NSCs (active or quiescent), since a large number of stem cells may promote differentiation.

In the following, we focus on a few scenarios, based on the biologically-motivated reasoning and having confirmed their good fit (Fig. 3). These are described by Eq. (3)-(7) and their fits are shown in Fig. 4A, with the respective dynamics of their system parameters (Fig. 4B). One important aspect to be noted is related to a bifurcation that appears at the point where the parameter *b*_0_ = 1*/*2 (see mathematical analysis in Methods). In particular, if *b*_0_ ≤ 1*/*2 the system only has the trivial steady state, whereas if *b>* 1*/*2 there exists a stable positive steady state. Depending on the regulatory feedbacks considered, upon parameter estimation, we obtain cases of good fits in which the system slowly converges to zero, and cases in which the system converges to the positive steady state (Fig. 4A). Interestingly, we observe that the estimations for scenarios in which qNSCs are also involved in regulating self-renewal (Fig. 4A, Eq. (4),(6)) lead to dynamics with positive steady state.

**Figure 4:**
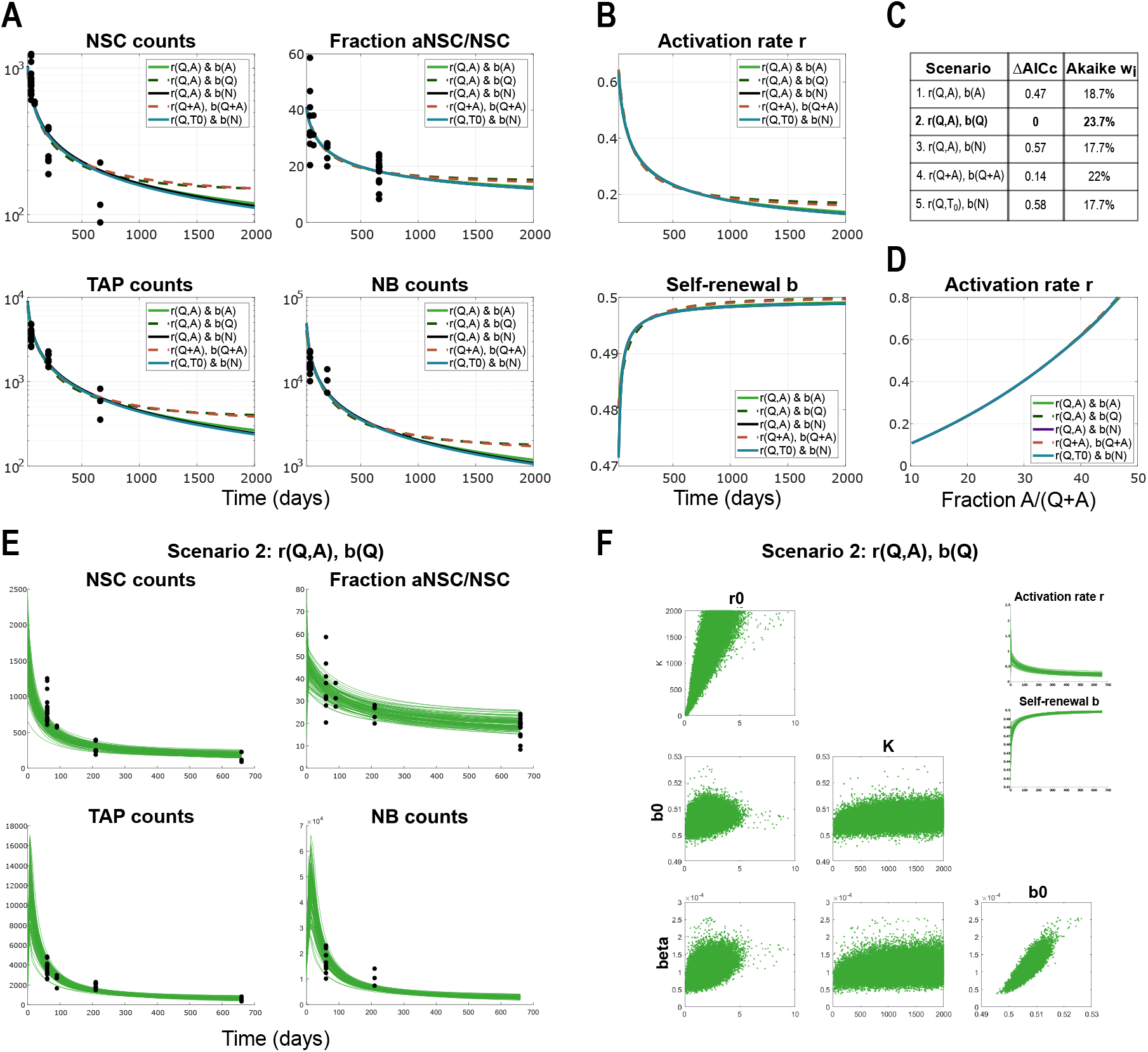
Model results and comparison among the five scenarios considered, applied to data from WT mice: **A**. Results showing the dynamics of lineage cell subpopulations in time, from the parameter estimation for five di erent scenarios in which the system parameters are regulated by lineage cell populations. Here NSC counts represent *Q*(*t*) + *A*(*t*), fraction of active corresponds to *A*(*t*)*/*(*Q*(*t*) + *A*(*t*)), TAP counts is considered the sum 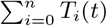 andNB counts is given by *N* (*t*). The scenarios considered in these plots are described by equations (3)-(7). Solid lines represent scenarios in which the estimated parameter *b*_0_ *<* 0.5 and the system only admits the trivial equilibrium. Dotted lines correspond to scenarios in which *b*_0_ *>* 0.5 and thus a stable positive steady state exists. X-axis represents Time, i.e. the age of mice in days. Y-axis is in logarithmic scale in all plots except for the one depicting the fraction of active cells. For readability, in the plot legends we write the multivariate functions *r*(*Q* + *A*) := *r*(*Q* + *A, Q* + *A*). **B**. Dynamics in time of the system parameters, activation rate *r* and fraction of self-renewal *b*, from the ve scenarios considered in panel A, described by equations (3)-(7). C. Model comparison based on AICc values and their respective Akaike weights. The best-scoring model according to Akaike metrics is Scenario 2, Eq. (4) with *r*(*Q, A*), *b*(*Q*). D. Activation rate *r* plotted against the fraction of active NSCs *A/*(*Q* + *A*) for the ve scenarios, showing an almost linear correlation. X-axis is *A/*(*Q* + *A*), y-axis is *r*. **E**. Trajectories of the solutions to Scenario 2, *r*(*Q, A*), *b*(*Q*) (Eq. (4)) corresponding to initial conditions sampled from a Gaussian distribution with mean and variance extrapolated from the available data, and parameter estimates from 250000 MCMC chain samples. The Y-axis is linear scale. **F**. Posterior distributions of model parameters and their correlation plots from the 250000 MCMC chain samples. Right-hand side insets show the time-course of system feedback functions *r* and *b* computed for the selected parameter values from the MCMC chain.

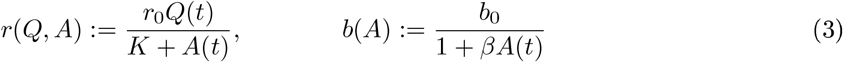

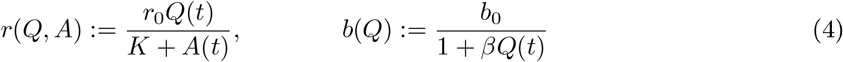

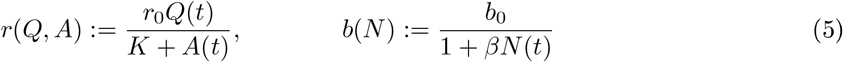

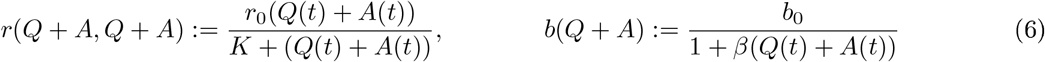

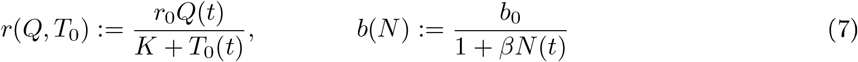

To confirm the validity of our parameter estimation methods and results, we quantify the sensitivity of the model to initial conditions and parameters by employing Bayesian methods. First, by assuming a Gaussian distribution of initial conditions with mean and variance derived from the available data points (please see Methods), we observe that all solution trajectories converge to the originally obtained time dynamics (Supplementary Figure A), indicating robustness. This supports our choice to fix initial conditions to reduce the dimensionality of the parameter space (see Methods). Furthermore, employing the Adaptive Metropolis algorithm 41 to generate Markov Chain Monte Carlo (MCMC) chain samples for our different model choices, we obtain posterior distributions for the model parameters that yield highly consistent trajectories (Fig. 4E). We observe a wide distribution of parameter *K*, which means a little influence on this parameter on the system feedback function. Therefore, this parameter cannot be confidently estimated from the available population dynamics data. Because of the high variability of *K*, we manually limited its range during MCMC sampling. The plots show also that the key parameters of the activation and self-renewal, *r*_0_ and *b*_0_ respectively, are weakly correlated, while *b*_0_ and *β* are strongly correlated, what can be well explained by the form of the feedback functions. Moreover, the feedback functions *r* and *b* that govern the dynamics of the solution exhibit no significant variation across samples. These results suggest a certain robustness of the neural stem cell system. Importantly, our Bayesian analysis shows that parameters from posterior distributions give comparable goodness of fit levels to the parameter estimates obtained before by the deterministic approach.

With the aim of gaining insight into which cell populations are involved in regulating the dynamics of the neural lineage, we compared the model scenarios under consideration. The most prominent difference among these scenarios emerges between the two groups of trajectories (with *b*_0_ ≤ 1*/*2 and *b*_0_ *>* 1*/*2) converging to different steady states (Fig. 4A). However, due to the limited time span of the available data and the fact that the divergence between these groups occurs later in the time course, beyond the typical lifespan of mice, we cannot determine a priori which case should be excluded from a biological standpoint. This underscores the need for targeted experiments that could address this aspect and inform model selection. A comparison of the AICc scores across the five scenarios (Fig. 4C) suggests that the hypothesis in which qNSCs promote and aNSCs inhibit activation, while self-renewal is negatively regulated by qNSCs (Scenario, Eq. (4), “*r*(*Q, A*), *b*(*Q*)”) is the best-scoring. However, its Akaike weight of 3.7% indicates only modest evidence, with Scenario 4, Eq. (6), in which the total NSC population is involved in both promoting activation and inhibiting self-renewal (“*r*(*Q* + *A, Q* + *A*), *b*(*Q* + *A*)”) following closely at %. The other scenarios cannot be easily discarded either, with their Akaike weights of 17.7-18.7%. Mathematical analysis further confirms that all five scenarios exhibit qualitatively similar solution behaviour (see Methods).

Given the similarity in the ability of the different scenarios to capture the dynamics observed in the data (Fig. 4A-B), we resorted to using more advanced optimisation methods and model discrimination techniques for optimisation-based model validation. Specifically, we applied the Multiple Shooting method in combination with the Gauss-Newton method 4, 43 to improve parameter estimation and model selection. These methods were used to assess whether a better fit could be obtained for some hypotheses, with the goal of distinguishing among well-fitting scenarios. Since the two parameter estimation approaches yielded nearly identical parameter values and fits, we performed F-tests to identify false models and to determine whether some models were still significantly different. When these tests were inconclusive, we used an optimal experimental design to explore the experimental conditions necessary to maximise the discrepancy among alternative scenarios.

Since discriminating among these hypotheses solely based on WT data was not possible, we supplemented our investigations with data from experiments in which the biological system was perturbed. In this work, we consider two biological perturbation settings, namely 1) killing actively dividing lineage cells with the chemotherapeutic drug Temo olomide (TM) 1 and) genetically knocking out the receptors of the inflammatory signals interferon *α* − *γ* (IFNAGR KO) 36.

### 2.4. Model validation based on perturbation experiments

#### 2.4.1. Model of TMZ treatment suggests regulation of the system by the entire NSC population and describes feature selection upon injury

Temo olomide (TM) is a chemotherapeutic drug that can be used experimentally to kill actively proliferating cells, which in our system comprise of aNSCs (*A*) and TAPs (*T*_*i*_). The authors of 1 performed experiments perturbing the system with TMZ in young (2-months old) and old (22-months old) mice, and recorded the recovery trend at 1, 9 and 35 days post-treatment. It was shown that in old mice the recovery is not as efficient as in young mice, so that either the numbers of proliferating cells do not recover back to the values pre-treatment, or it is much slower such that it is not seen in the first 35 days post-treatment. To gain insights into the mechanisms of recovery and the differences between young and old mice, we applied the various scenarios of the nonlinear model to the available data from the TMZ-perturbed system.

By performing the TMZ experiments in silica with the parameters estimated in WT data, and plotting the results we observe that the recovery is too slow compared to the data from young mice and too quick compared to that from old mice (Supplementary Figure 1B, left). Even when trying to perform parameter estimation to fit all WT data and young and old TMZ data together with the same set of parameters (potentially allowing a slightly worse fit of WT data than before), the results are not satisfactory (Supplementary Figure 1B, right). We can thus conclude that perturbing the biological system by administering the TMZ drug changes not only the behaviour of system parameters (*r* and *b* functions) but values of individual model parameters as well.

Next, in the attempt to maintain the view as general as possible, we checked whether we can fit the data from TMZ-perturbed young and old mice with the same parameter set, but different from that estimated for WT. Intuitively, even if we keep the same model parameters in both cases, the recovery could be different due to the regulatory feedbacks among cell populations of different sizes. This is however also not the case, suggesting that not only does TMZ change the properties (parameters) of the cells but it does it differently when administered at different ages. Thus, the parameter estimation should be performed separately for young and old mice.

Proceeding step by step, we first assumed that the proliferation rates *p*_*A*_ and *p*_*T*_ were the same as in WT and not influenced by TMZ treatment, and only allowed model parameters in *r* and *b* to change. Even though the data itself could be well fitted, the solution converged to a different steady state that was high above the one from the healthy non-perturbed WT data (Supplementary Figure 1B, middle). This seems unreasonable as it would mean that applying a chemotherapeutic drug that kills cells makes the biological system performing much better long-term. Even though this could be the case short-term, one would not expect that such a strong aggressive perturbation would prevent ageing in the long run. This result suggests that either TMZ treatment changes the parameters for a short period of time and later the system switches or transitions back to the WT parameters, or there is a different process at play, one that allows the recovery after death as well as a return to the WT behaviour after some time, whilst keeping the parameters that were estimated based on the recovery trend data. The latter was further investigated by also allowing *p*_*A*_ and *p*_*T*_ to change upon TMZ treatment.

Once again, by trying to keep the model as simple as possible and the changes due to TMZ treatment as few as possible, in order to find the essential aspect that changes upon treatment, we took a stepwise approach by allowing as few parameters as possible to differ from those from WT. We found that in young mice all parameters can remain identical as in WT apart from the proliferation rate of TAPs, *p*_*T*_, the estimated value of which is reduced by approximately 60% compared to untreated WT mice. This means that TMZ treatment affects TAPs such that they proliferate much slower than before (lengthening of cell cycle duration). Considering a heterogeneous population of proliferating cells with respect to their cell cycle length, our model parameter *p*_*T*_ represents a mean value of the heterogeneous proliferation rates. In this context, our results suggest that TMZ kills fast proliferating TAPs and thus select for slow dividing ones, which corresponds to the decreased value of *p*_*T*_ after treatment in the model. Nevertheless, properties of aNSCs seem to be unaffected by the perturbation. As far as old mice are concerned, TMZ treatment also reduces *p*_*T*_ (by approximately 80%), and in addition the *r*_0_ parameter is decreased by one order of magnitude, which means that NSCs become arrested into a deeper state of quiescence, leading to a smaller mean *r*_0_ of qNSCs. This is also in agreement with previous results 12] that stated that quiescent NSCs in old mice are more resistant to injury-induced activation. Additionally, we find that proliferating cells of young mice are more resistant to TMZ treatment than old mice, with a death rate of approximately 1.5-fold difference upon drug administration. With these assumptions, the model is able to recover the dynamics observed in the data (Fig. 5A-A’). We observe that some neural populations are increased post-treatment and their dynamics remain above those of the unperturbed system, while the opposite is true for others (Fig. 5B-C’). In young mice, the population of TAPs after recovery is bigger than that in WT and this is due to the slower proliferation rate *p*_*T*_, which eventually leads to a slower differentiation into NB and thus less exit from the neural system (Fig. 5B bottom). In old mice, the dominant parameter post-treatment is *r*_0_, the decrease of which preserves a more quiescent population and, together with the lower value of *p*_*T*_, leads to a lower recovery of proliferating and differentiated cells (Fig. 5B’, bottom).

**Figure 5:**
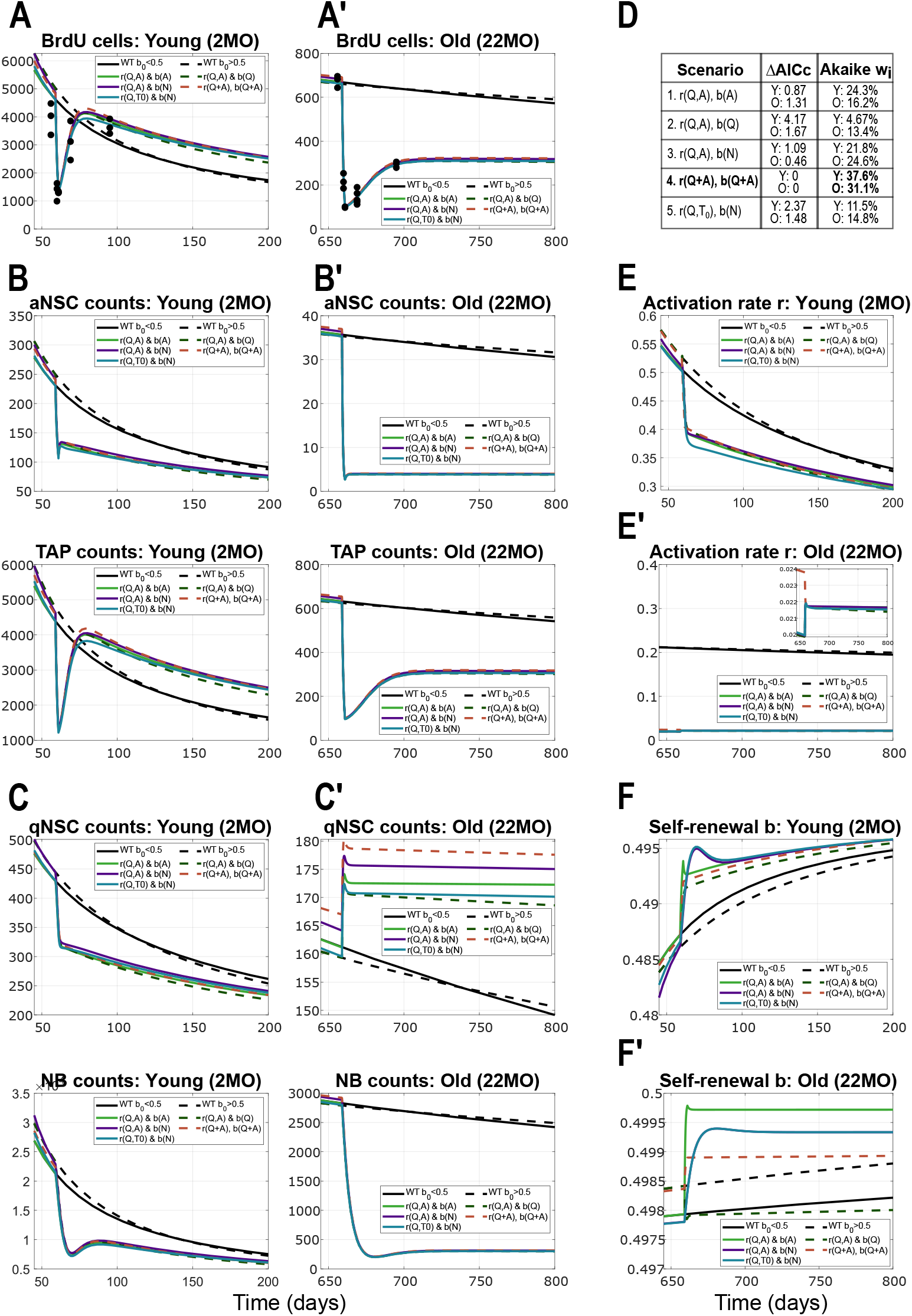
Model results and comparison among the five scenarios considered, applied to data from TMZ treatment. Panels are split into young (letter X) and old (letter X’). **A-A’**. Results showing the recovery of proliferating cells (BrdU positive, consisting of aNSCs and TAPs) after TMZ treatment, in young (2MO) and old (22MO) mice. Throughout Fig. 4, black (solid and dotted) lines correspond to 2 scenarios of the WT unperturbed dynamics. More specifically, the two WT scenarios depicted are: *r*(*Q, A*), *b*(*A*) (solid line) and *r*(*Q, A*), *b*(*Q*) (dotted line). Solid lines (both WT and perturbed) correspond to models where *b*_0_ *<* 1*/*2 (with trivial steady state), and dotted lines to scenarios in which *b*_0_ *>* 1*/*2 (having a stable positive steady state). **B-B’**. Dynamics of aNSCs (top) and TAPs (bottom), which together form the BrdU cells from panel A, in young (B) and old (B’) mice. **C-C’**. Dynamics of the populations of qNSCs (top) and NBs (bottom) in time, as a result of TMZ intervention on BrdU cells, in young (C) and old (C’) mice. **D**. Table comparing the five scenarios based on AICc values, and their Akaike weights quantifying the individual probabilities for selection. The model in which the entire population of NSCs (*Q* + *A*) regulates both *r* and *b* scores the highest (Scenario 4, Eq. (6), “*r*(*Q* + *A, Q* + *A*), *b*(*Q* + *A*)”). **E-E’**. Activation rates *r* in the case of TMZ treatment compared to the WT setting, in young (E) and old (E’) mice. **F-F’**. Fraction of self-renewal *b* upon TMZ treatment in comparison to the case of the unperturbed WT setting, in young (F) and old (F’) mice. For readability, in the plot legends we write the multivariate functions *r*(*Q* + *A*) := *r*(*Q* + *A, Q* + *A*).

Aiming at selecting a model scenario with the help of the additional TMZ perturbation insights, we computed the Akaike scores and weights as before. In this case, the best-scoring scenario was that in which the system parameters are both regulated by the total population of NSC, Eq. (6), with Akaike weights of 3 .6 for young mice and 3. for old mice (Fig. 5). Surprisingly, the best-scoring scenario found for the WT data scored the lowest in the case of the TMZ perturbation. Furthermore, in an attempt to better discriminate among models, we applied optimal experimental design methods to determine the age of mice to receive TMZ treatment in order to have as much discrepancy between scenarios as possible. We found that the optimal treatment age was in the interval 60,500] days old and again suggested that having available measurements of the qNSC subpopulation after treatment would increase our confidence of the model selection results, as this compartment showed the greatest differences across scenarios.

Overall, the modelling of TMZ experiments has provided interesting insights into the behaviour of the neural lineage system and the changes that appear due to treatment, and suggested that upon injury the system parameters are regulated by the entire NSC population.

#### 2.4.3. Model of IFNAGR Knock-Out stays in agreement with insights from wild-type and describes involvement of non-neural cells in modulating activation

Proceeding further with our plan to uncover the most likely regulatory feedbacks, we decided to re-tackle the IFNAGR KO perturbation from [36]. A closer look at the data on the fractions of active NSC among all NSC (Fig. 6A, top right), suggests that in the KO mice the values grow with ageing. Additionally, we observe that generally the values of these fractions are very well, almost linearly, correlated to the activation rate (Fig. 4E), which implies that the activation rate in KO data also increases in time. Due to the way in which the time-dependent *r* and *b* functions were defined in 36], the linear non-autonomous models were not able to capture the slight increase in the fraction of active NSC.

**Figure 6:**
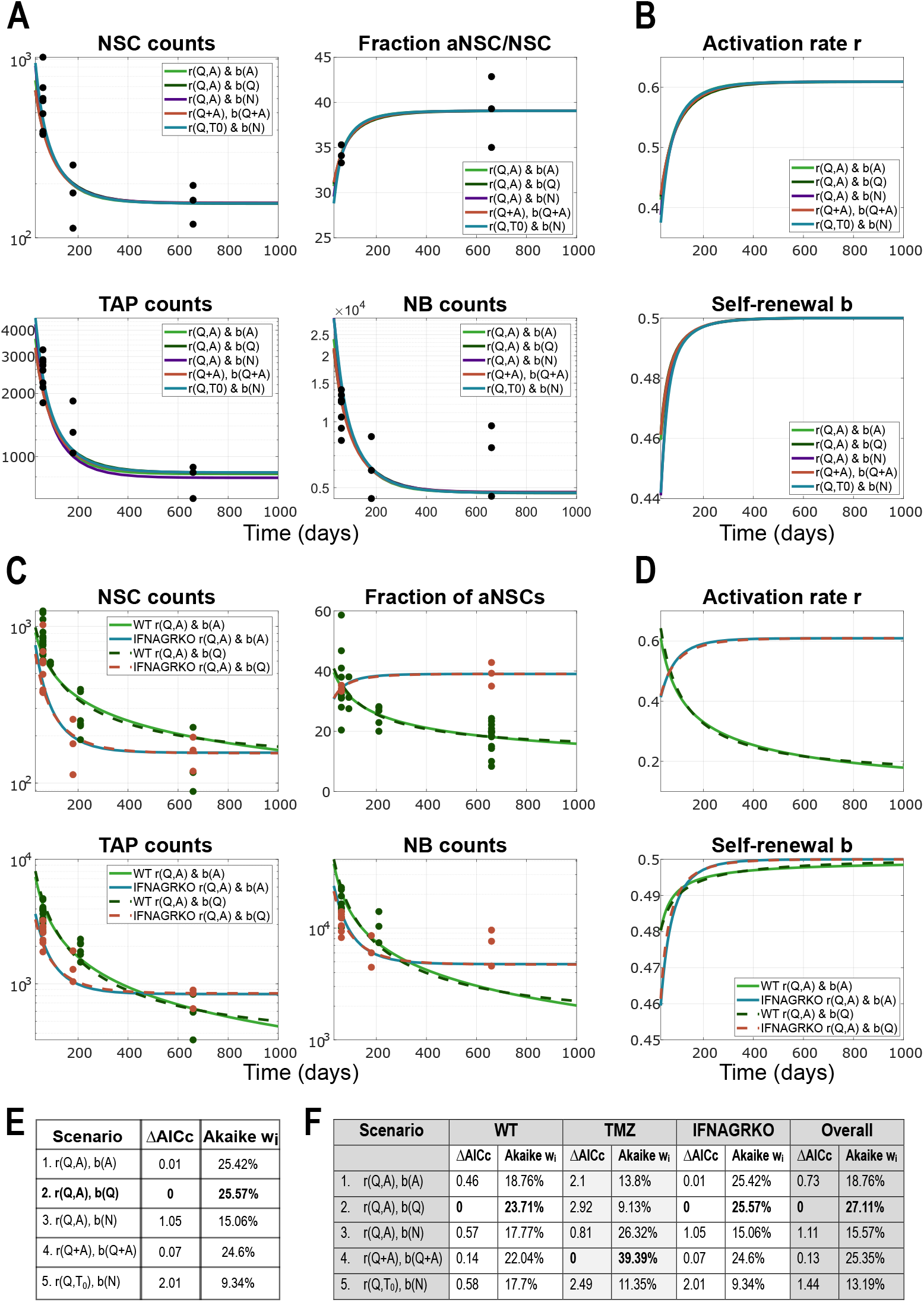
Model results and comparison among the five scenarios considered, applied to data from IFNAGR KO mice: **A**. Simulation results showing the dynamics of lineage cell subpopulations in time, from fitting to the IFNAGR KO data, for the five different scenarios. The scenarios considered in these plots are given by equations (9)-(13). Parameter estimation finds *b*_0_ *>* 0.5 in all five scenarios and thus the system has a stable positive steady state to which it converges. X-axis represents Time, i.e. the age of mice in days. Y-axis is in logarithmic scale in all plots except the one depicting the fraction of active cells. **B**. Dynamics in time of the system parameters, activation rate *r* and fraction of self-renewal *b*, from the five scenarios considered in panel A. **C**. Comparison between WT and IFNAGR KO fits to their respective data for two scenarios: *r*(*Q, A*), *b*(*A*) (solid lines, for WT *b*_0_ *<* 0.5) and *r*(*Q, A*), *b*(*Q*) (dotted lines, for WT *b*_0_ *>* 0.5). **D**. Dynamics of the activation rate and fraction self-renewal for comparing IFNAGR KO with WT, for the scenarios considered in C. **E**. Table with AICc values for model selection and their respective Akaike weights, for the model of IFNAGR KO dynamics. **F**. Overview table with AICc scores and Akaike weights for each scenario, computed separately for WT, TMZ and IFNAGR KO, as well as combined for all settings and data together (“OverallW). For readability, in the plot legends we write the multivariate functions *r*(*Q* + *A*) := *r*(*Q* + *A, Q* + *A*).

Using the nonlinear models with population-dependent *r* and *b*, we infer that, in order for *r* to increase in time, we need a slight decrease with respect to cell counts. This hints towards an inhibitory-type Hill function for *r* in the case of IFNAGR KO mice, similar to that for the self-renewal parameter *b*. Therefore, in order to allow capturing the time evolution of neural subpopulations from both WT and KO mice with the same hypothesis, we assume the following function for *r* whilst keeping the *b* function as before.

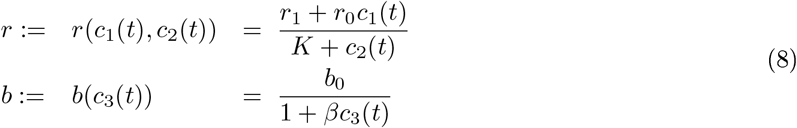

As a result, our five scenarios from Eq. (3)-(7) are extended for the IFNAGR KO setting to Eq. (9)-(3).

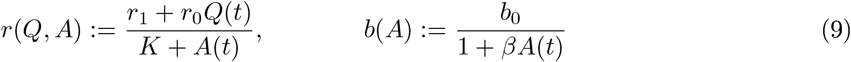

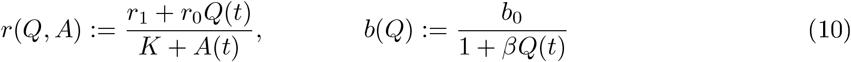

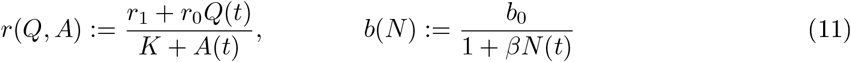

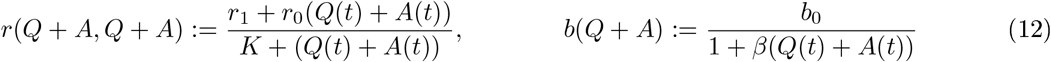

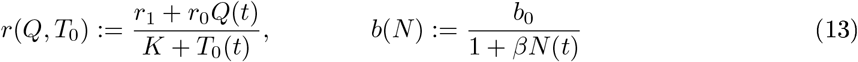

The simulation results for these scenarios are depicted in Fig. 6A and show that all model scenarios are very well able to capture the trend of the data. As expected based on the construction as in Eq. (8), both the activation rate *r* and the fraction of self-renewal *b* increase in time (Fig. 6B). Depending on the different estimated values of *r* parameters (i.e. *r*_0_, *r*_1_ and *K*), this definition of the *r* function (Eq. (8)) allows to capture both the decreasing fraction of active cells in WT and the increasing one in KO mice. The previous activation mechanisms modeled in the WT setting by the activating Hill-type function (Eq. (2)) is a particular case of the model with system parameters given by Eq. (8), with *r*_1_ = 0. The posterior distributions obtained from the MCMC chains simulated with the Adaptive Metropolis algorithm, further emphasize the importance of the *r*_1_ ≠ 0 parameter in the IFNAGR KO experiment (Supplementary Figure 2B-C). An additional insight that we can draw from the comparison between WT and IFNAGR KO results (Fig. 6C-D), is that in KO mice the activation is disregulated and consequently the self-renewal compensates (Fig. 6C-D), acting as a secondary layer of regulation, as also previously suggested [36]. Because a higher activation rate that does not decrease with ageing can lead to a fast depletion of the NSC pool, the self-renewal has a more drastic increase to counteract it. Additionally, parameter estimation suggests that the proliferation rate of TAPs is slightly higher than that in WT mice, corresponding to a cell cycle length of approximately 16h (parameter values in Supplementary Table 1).

Once again, Scenario 2 (Eq. (10)) is found as best-scoring by the Akaike model comparison framework, closely followed by Scenarios 1 (Eq. (9)) and 4 (Eq. (12)), similar to the the WT setting. Mathematical modelling thus allows us to gain new insights into the dynamics of neural lineage cells and into the dialogue bewteen neuroinflamation and adult neurogenesis, despite not clearly uncovering how the regulation is performed.

### 2.5. Interpretation in the context of signalling molecules

The dynamics of the system parameters *r* and *b* arise from a complex network of signalling pathways acting both within and beyond the neural niche. By analysing how these parameters depend on various system components, we can begin to identify which cells produce or respond to signalling molecules that regulate activation and influence the balance between self-renewal and differentiation. To explore this mechanistically, we extend the feedback model (Eq. (8)) to explicitly represent the dynamics of the signalling molecules mediating feedback. This approach-essentially a reverse quasi-steady-state reduction-is inherently challenging, as the observed regulatory effects may result from the combined action of multiple signals.

We may capture this complexity in a tractable form by introducing two effective signals, *S*_*r*_ and *S*_*b*_, which serve as representative examples of the underlying regulatory mechanisms. These abstract variables substitute for the parameters *r* and *b*, and their dynamics are governed by additional differential equations (Eq. (14)) that account for signal production, degradation, and internalisation. Terms associated with cell types *c*_*i*_ describe their contribution to the signal pool—either as producers or consumers—depending on the sign of the corresponding term. Basal production from non-lineage supporting cells is captured by positive constants, while degradation is modelled through negative linear terms.

Assuming fast signal kinetics, we the extended model can be reduced back to the nonlinear feedback formulation with Hill-type regulation. In this way, the effective signals *S*_*r*_ and *S*_*b*_ illustrate how abstracted feedback terms may reflect more complex, mechanistically grounded regulatory interactions. (see [44, 45]):

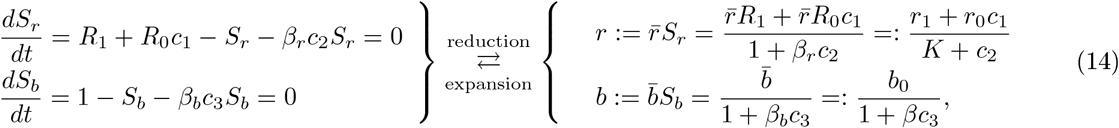

with 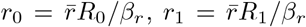, *K* = 1*/β*_*r*_, 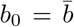 and *β*_*b*_ = *β*, to recover the form and notation from before.

Consequently, this model extension provides a mechanistic interpretation of the previously inferred Hill-type feedback functions in terms of specific signalling processes. Based on this generic formulation, we deduce that, in wild-type (WT) mice, the signal promoting NSC activation is both produced and internalised by neural lineage subpopulations-potentially distinct-corresponding to the *c*_1_ and *c*_2_ cell types in Eq. (2), respectively. In this context, we set *r*_1_ = 0, assuming no contribution from non-neural sources to activation. In contrast, the signal controlling the balance between self-renewal and differentiation of aNSCs appears to be produced by non-neural supporting cells and consumed by neural lineage cells (type *c*_3_). These findings align with existing experimental evidence for autocrine and paracrine signalling. For instance, canonical Wnt signalling-secreted by astrocytes and vascular endothelial cells and taken up by neural cells-is believed to play a central role in promoting aNSC self-renewal.

When applying the nonlinear model to experimental data from Ifnar1/Ifngr1 double knockout (IFNAGR KO) mice, we observe that signal production by neural populations is impaired, rendering their contribution to activation negligible (*r*_0_ ≈ 0). As a result, non-neural supporting cells assume a significantly greater regulatory role in driving system dynamics.

More complex alternative mechanistic scenarios may also give rise to Hill-type regulatory functions, providing complementary interpretations that converge on similar forms of system-level regulation. While several signalling pathways have been identified as inhibitors of activation, comparatively little is known about signals that actively promote it. For instance, Notch and Wnt signalling are well-established regulators of adult neurogenesis and specifically influence key system parameters such as activation rate *r* and self-renewal fraction *b*. It has been shown that aNSCs express Delta ligands, which bind to Notch receptors on neighbouring qNSCs, thereby suppressing their activation. Assuming an inverse relationship between the activation rate *r* and the Delta signal and applying a quasi-steady-state approximation to a dynamical model (ODE) describing Delta dynamics naturally yields a Hill-type function for *r*, see Eq.15. This formal derivation provides a plausible mechanistic underpinning for the empirically inferred feedback structure and illustrates how inhibitory autocrine or juxtacrine signalling can manifest as effective population-level regulation.

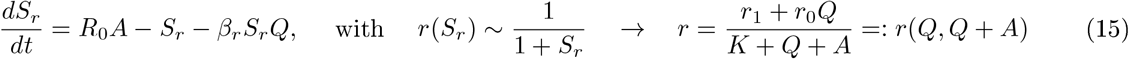

Furthermore, Wnt signalling-another key regulator of adult neurogenesis-is predominantly secreted by non-neural niche components, such as astrocytes and endothelial cells, and exerts its effects primarily on quiescent NSCs. This functional specificity is consistent with the inferred shape of the self-renewal parameter *b*(*Q*) in Scenario 2, Eq. (10). Taking into account these regulatory processes leads to a new scenario, “*r*(*Q, Q* + *A*), *b*(*Q*)” (Eq. (16)), which is in fact a combination of our two best-scoring hypotheses (Fig. 6F), offering a biologically grounded and mechanistically coherent explanation for the observed dynamics.

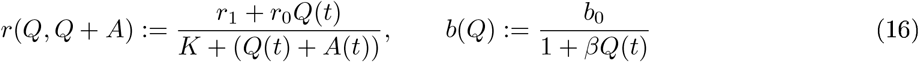

A graphical comparison between the additional scenario and the two previously best-scoring models is shown in Fig. 7A-B. MCMC simulations of the new scenario revealed that, for data from WT mice, the parameter *r*_1_, originally introduced to model the IFNAGR KO condition (Eq. (8); Scenarios in Eq. (9)-(13)) and also used in the Delta-Notch hypothesis (Eq. (16)), plays no substantial role, thereby supporting our original model choice (Eq. (2)). In contrast, for the IFNAGR KO data, the parameter *r*_1_ /= 0 emerges as the most influential parameter, overtaking *r*_0_. While models for WT data (with *r*_1_ = 0, Eq. (3)-(7)) exhibited a strong correlation between *r*_0_ and *K* (Fig. 4F), this correlation shifts to *r*_1_ and *K* in the extended models for the IFNAGR KO condition (Eq. (9)-(13)) and the Delta-Notch-Wnt hypothesis (Eq. (16)). Notably, including the additional parameter *r*_1_ in WT models increases uncertainty and reduces identifiability, with no critical impact on the fit to the data. These observations are consistent across all scenarios. Fig. 7D illustrates how the Delta-Notch-Wnt hypothesis captures the recovery following TMZ treatment in both young and aged mice (for simplicity, *r*_1_ was set to zero). When comparing AICc values and Akaike weights for the additional scenario against the two previously best-scoring ones, we find all three to be equally plausible in terms of explaining both WT and IFNAGR KO data. Consistent with previous observations (Fig. 5D, 6F), analysis of the TMZ-perturbed data favors the scenario in which the total NSC population regulates both activation *r* and self-renewal *b*, with high confidence (Fig. 7C).

**Figure 7:**
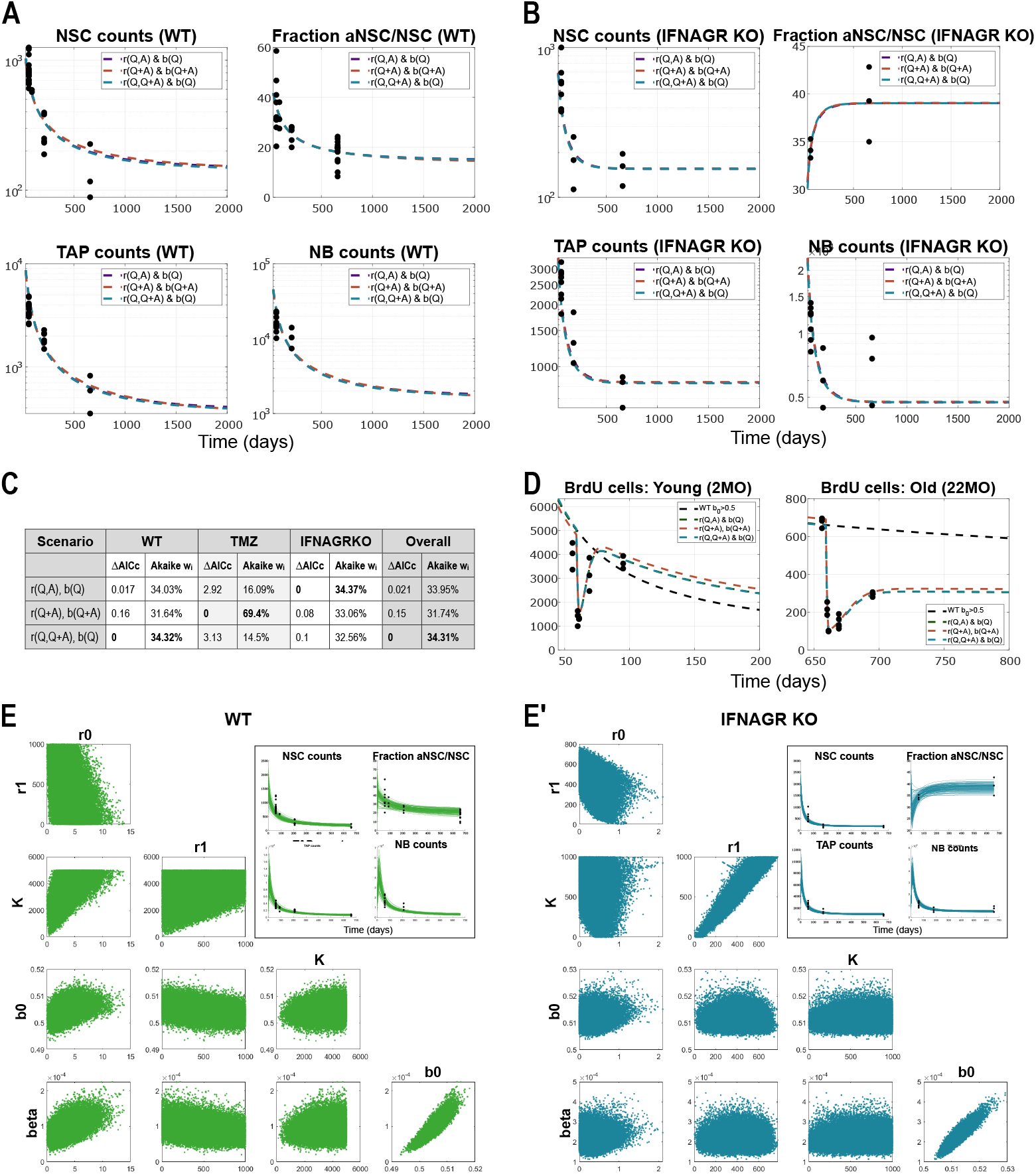
Model results and comparison among the two best-scoring scenarios and the Delta-Notch-Wnt scenario “*r*(*Q, Q* + *A*), *b*(*Q*)”, applied to data from WT and IFNAGR KO mice: **A**. Simulation results showing the dynamics of lineage cell subpopulations in time, from fitting to the WT data, for the three different scenarios. The scenarios considered in these plots are given by equations (10),(12) and (16). Parameter estimation finds *b*_0_ *>* 0.5 in all three scenarios and thus the system has a stable positive steady state to which it converges. X-axis represents Time, i.e. the age of mice in days. Y-axis is in logarithmic scale in all plots except the one depicting the fraction of active cells. For readability, in the plot legends we write the multivariate functions *r*(*Q* + *A*) := *r*(*Q* + *A, Q* + *A*). **B**. Simulation results showing the dynamics of lineage cell subpopulations in time, from fitting to the IFNAGR KO data, for the three different scenarios. **C**. Table with AICc values for model selection and their respective Akaike weights, for the three model scenarios depicted in A-B. **D**. Simulation results of the three scenarios showing the recovery of the population of BrdU cells after treatment with TMZ, in young (left) and old (right) mice. **E-E’**. Plots of posterior distributions of model parameters and their correlation plots from the samples of the MCMC chain, in WT (E) and IFNAGR KO (E’) mice. The insets show the dynamics of the solutions corresponding to initial conditions sampled from a Gaussian distribution with mean and variance extrapolated from the available data points, and parameters sampled from their posteriors.

Taken together, these considerations culminate in a biologically plausible feedback scenario grounded in Delta-Notch and Wnt signalling, which we propose as a mechanistic explanation for the observed regulation of activation and self-renewal. However, while this scenario captures key aspects of the data and aligns with known biology, we acknowledge that additional, possibly synergistic, signalling molecules may contribute to the robustness and adaptability of the system. Indeed, the effective signalling model (Eq. (14)) could be extended to encompass a broader network of interacting signals. For instance, the activation rate could be modulated by two distinct signalling components: a promoting signal *S*_*rp*_ and an inhibitory signal *S*_*ri*_. In this extended framework, *S*_*rp*_ would be produced by neural lineage cells (analogous to *c*_1_) and degraded either spontaneously or by non-lineage cells, whereas *S*_*ri*_ would be secreted by non-neural supporting cells and received by neural lineage cells (as *c*_2_). Alternatively, similar regulatory effects could arise through a double-inhibition mechanism, in which an inhibitory signal is itself suppressed by lineage cells-mirroring the Delta-Notch regulation of activation. These more complex signal interactions illustrate that the feedback architecture inferred here may reflect an aggregate effect of multiple pathways. Elucidating the precise molecular components and their interactions will require targeted experimental validation and lies beyond the scope of the present study.

## 3. DISCUSSION

In this paper, we aimed at deciphering regulatory feedbacks acting within the neural lineage, which would be leading to the observed age-related changes in adult neurogenesis in the ventricular-subventricular zone of adult mice brains. Even though data of healthy neurogenesis from humans are not available, recent work studying the architecture and progression of the glioblastoma brain cancer showed on one hand, that patient tumour samples comprise the entire neural lineage as in healthy mice, and on the other hand that tumour samples transplanted into mouse brains show the same properties in the host as in the donor [46]. These considerations hint to a good agreement between human and mouse neurogenic systems. We suggested feedback mechanisms that replicate the observed dynamics and which are also plausible from a biological perspective. We additionally derived a minimal scenario based on biological evidence of signalling among NSCs via Delta-Notch and Wnt pathways. Although model selection did not prove entirely conclusive, we found a number of insights that can guide further experiments and investigations. First of all, we showed that the decision of a neural stem cell to self-renew or differentiate is negatively regulated by the neural lineage. From the perspective of signalling, self-renewal signalling molecules are produced by supporting cells and bound by lineage cells. Additionally, we found that the balance between activation and quiescence is mainly regulated by the neural populations, not unlikely with the involvement of two different subpopulations acting in opposing directions. The mathematical models presented a few ways through which quiescent stem cells promote their activation in terms of signalling mechanisms. Considering the intricacy of the network of signalling interactions involved in regulating the system parameters in adult neurogenesis, it is not easy to single out specific genes, but modelling can shed light onto how and where the regulation acts.

In addition, we highlighted the risks of inference from preliminary models applied to inconclusive or insufficiently informative data without conducting a thorough and unbiased investigation, as multiple hypotheses may lead to similar results despite differing interpretations. Further, to increase the power of model selection, we used supporting data from two different perturbation experiments: the treatment with the chemotherapeutic drug TMZ, and the knock-out of the IFNAG receptors. The mathematical models applied to these complementary data uncovered compelling insights. We showed that upon TMZ treatment, the mean proliferation rate of TAPs is drastically reduced and we argue that it is likely a consequence of feature selection in a heterogeneous population: fast-proliferating TAPs are killed by TMZ and the ones that escape are those with longer cell cycle, hence lower proliferation rate. A similar reasoning applies to old mice, where in addition the maximum activation rate is greatly reduced, leading to a surviving population of more deeply quiescent stem cells. We observed that due to slower proliferation of TAPs but a steady influx from the active NSC compartment, the total population of TAPs decays more slowly after TMZ treatment in young mice, and that this leads to a trend that lies higher than in the unperturbed system during the lifespan of mice (Fig. 5B, bottom). However, the populations of stem cells remain smaller than before treatment (Fig. 5B-B’, top). Interestingly, the subpopulation with the most dissimilar behaviour across scenarios is that of quiescent NSCs (Fig. C-C 1 top). Although the dynamics is similar1 the values of cell counts differ. Would it be possible to better select the most plausible model hypothesis if data on numbers of quiescent NSCs after treatment were available? Such data quantifying qNSCs might improve model selection assuming the heterogeneity among mice is not too high. Ideally1 an additional experiment could focus on only perturbing either active NSCs or TAPs1 which might help uncover whether feedback originates solely from the NSC compartments or also from more differentiated neural subpopulations. Furthermore1 perturbation of the Delta-Notch pathway would provide new data which together with our models could restrict our set of hypotheses and offer new insights into the regulatory mechanisms.

With respect to the perturbation by knocking out IFNAG receptors1 considering that data show an ascending trend of the fraction of active NSCs among all NSCs1 we showed that the activation rate of quiescent NSCs increases with ageing. From the perspective of signalling1 we suggested that in the IFNAGR KO mice the activation signal is no longer produced by neural cells1 but by supporting cells. Additionally1 as the activation of quiescent NSCs is disregulated in IFNAGR KO mice1 self-renewal compensates via an earlier and faster increase. As activation is a major player in the recovery after TMZ treatment in WT mice1 it will be interesting to perform the TMZ experiment on IFNAGR KO mice and inspect how the neural cells behave.

Finally1 in terms of selecting a model hypothesis1 even though no definitive conclusion can be drawn1 observations can be made from simulation results. First of all1 parameter estimation for the various scenarios suggests that if qNSCs are involved in regulating the self-renewal1 the system asymptotically converges to a positive steady state. The nonlinear models suggest that the driving effect for exit from quiescence most likely originates primarily from qNSCs. To our knowledge1 experimental studies have gained insights into signalling mechanisms that inhibit activation1 but not much has been reported about promoters of activation. Does there exist a mechanism through which qNSCs actively drive their exit from quiescence1 or is the process solely regulated through a double-inhibitory feedback1 as derived for our Delta-Notch hypothesis? A new angle of experimental research could be aimed at uncovering potential mechanisms through which quiescent NSCs directly promote their activation. The table of model selection scores for the three individual settings (WT1 TMZ and IFNAGR KO) together with their overall score (Fig. 6F1 Fig. 7C) suggests that in an uninjured setting (WT or IFNAGR KO) the main regulator of self-renewal is the qNSC subpopulation1 whereas activation is primarily promoted and inhibited by qNSC and (a)NSC1 respectively (Scenario 21 Eq. (4)1(10) or the Delta-Notch scenario1 Eq. (16)). In the case of a severe injury such as the treatment with TMZ1 the entire NSC population is involved in regulating the system parameters to ensure a smooth recovery (Scenario 41 Eq. (6)). If we inquire how these scenarios influence the recovery1 we gather that having aNSCs contributing to regulating self-renewal (Scenario 41 (6)) leads to a faster increase in self-renewal upon their death by TMZ1 than when only qNSCs are involved (Scenarios given by Eq.(4) or (16))1 leading to a more efficient repopulation of the NSC pool. It is reasonable to assume that generally the two NSC subpopulations contribute with different weights in governing the system parameters1 however investigating this aspect by inserting additional parameters into the models is not feasible with the available data. Additionally1 our model selection results suggest that multiple neural lineage subpopulations might be involved in the regulations1 with various impact1 pertaining to an existing redundancy in the neural system that might ensure better adaptability. Altogether1 we find that the main regulator of the dynamics of the neural lineage is the population of NSCs1 whereas downstream subpopulations1 if involved1 have a much weaker influence.

## METHODS

The current work is based on the methodology of nonlinear ordinary differential equations and combines mathematical analysis1 computational simulations1 parameter estimation1 model selection and identifiability.

### Mathematical analysis

The crucial parameter describing the dynamics of the system (1) (Fig. 1B) is *b*_0_. It is clear that for *b*_0_ ≤ 1*/*2 it holds:

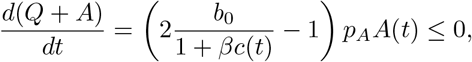

where *c*(*t*) can be equal to *A*(*t*), *Q*(*t*), *N* (*t*) or *Q*(*t*)+ *A*(*t*). It means that for all scenarios (3)-(7) the only steady state is a trivial one. Moreover, the solutions are asymptotically converging to it.

In the case when *b*_0_ *>* 1*/*2 there exist two steady states for each model: a trivial one and a positive one. For all scenarios (3)-(7) the trivial steady state (when *b*_0_ *>* 1*/*2) is unstable. The positive steady state is thus our focus in the rest of this section. We divide the models into two groups. The first comprises scenarios (3), (4) and (6), where parameters *r*(·) and *b*(·) depend only on *Q* and *A*. In these models, it is enough to consider the existence of the solution and its stability for the reduced system composed of the first two equations (1)_(1)_-(1)_(2)_, the results for the full model being a natural extension.

#### Theorem 1.

*There exists a unique g oba so ution to (1) with system parameters* (3), (4) *or* (6). *Moreover, for b*_0_ *>* 1*/*2 *the positive steady state solution is stable*.

*Proof*. The functions on the right-hand side of (1) are Lipschitz-continuous for nonnegative values of the solution. Starting with nonnegative initial conditions, we obtain the local existence of unique solutions to (1), according to the Picard-Lindelöf theorem. Using the property of the right-hand sides of (1) we obtain the positivity of the solutions.

To prove the global existence of the solution we focus on the system (1)_(1)_-(1)_(2)_. We will present here the results for the first scenario, i.e. (3). The reasoning in the other cases is similar. Then

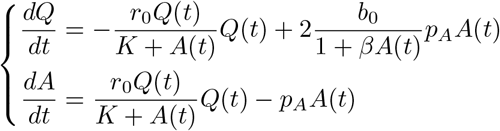

Equating the right-hand sides of the aforementioned equations to 0 we obtain two isoclines, which after transformation are equal to

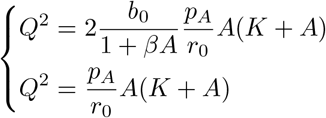

It is clear that for nonnegative values of *Q* and *A* these lines have at most two intersection points: one at (*Q, A*) = (0, 0), and a second one that occurs only when *b*_0_ *>* 1*/*2. The value of the second intersection point is equal to 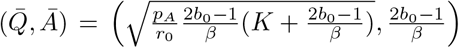. Note that if we decrease *b*_0_ to 1*/*2 from a greater value, then the second positive point goes to (0, 0).

We define a domain ℳ = {(*Q, A*) ∈ ℝ2 : *Q, A*<u>≥</u> 0 and *Q* ≤ *Q*^*\**^, *A* ≤ *A*^*\**^}, where *A*^*\**^ *> Ā* and *Q*^*\**^ is chosen such that 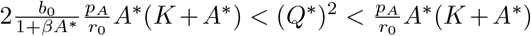. Then the solution with initial conditions in ℳ does not escape from ℳ. Thus the solution is global in time.

The existence of a global unique solution for the reduced system (1)_(1)_-(1)_(2)_ implies the existence of a global unique solution for the full (1) system, since *Q*(*t*) and *A*(*t*) are bounded everywhere.

The stability of the positive steady state is a consequence of the linearization of the model in the neighbourhood of the steady state.

□

We next focus on the second group of scenarios, where downstream populations are also involved in the regulation, i.e. (5) and (7).

#### Theorem 2.

*There exists a unique gobal soution to (1) with* (5) *or* (7). *Moreover, for b*_0_ *>* 1*/*2 *the positive steady state solution is stable*.

We will focus on the model with (5). Similar results can be obtained for the model with (7), but due to the higher dimensionality, we will omit the computations here.

*Proof*. Similarly to the proof of Theorem 1, we obtain the local existence of a unique positive solution as a consequence of Picard-Lindelöf theorem and properties of the right-hand side functions. It remains to define a domain ℳ from which the solutions cannot escape. We work with a reduced system containing only (*Q, A, N*), i.e. first two ODEs together with the last one in (1) further denoted (1 _*\**_). Results obtained for this model can be naturally extended to the full model (1). Moreover, for simplicity, we assume that *b*_0_ *<* 3*/*4.

The invariant domain ℳ for this model is a 3D polyhedral with 7 faces. The base in the (*A, Q*)-plane is quadrilateral. Two of their edges coincide with axes and the other two are defined by functions *Q* + *A* = *constant* and *Q − A* = *constant*. The upper base is larger than the bottom one (*N* = 0). The two faces that coincide with the planes *Q* = 0 and *Q* − *A* = *const* are pentagons. The other faces of the polyhedral are quadrilateral lying on the planes: *A* = 0, *Q* + *A* − *N* = *constant* and *Q* + *A* = *constant*.

We denote 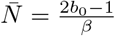. We take an arbitrary *<u>A</u>* greater than 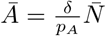. Moreover, *<u>Q</u>* and *N* ^*\**^ are chosen in the following way:

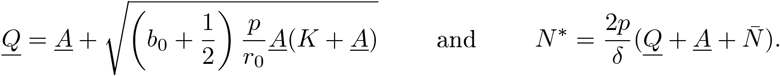

The coordinates of the vertices of the domain ℳ are as follows: {(0, 0, 0), (*<u>A</u>* + *<u>Q</u>*, 0, 0), (*<u>A</u>, <u>Q</u>*, 0), (0, *<u>Q</u>* −*<u>A</u>*, 0), (0, 0,*N* ^*\**^), 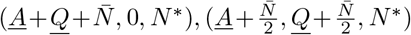, (0, *<u>Q</u>*−*<u>A</u>,N* ^*\**^), 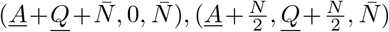} We need to prove that solutions starting from each of the seven faces of the polyhedral domain ℳ go inside the domain. We previously showed that our solutions are positive if the initial data are positive so the edges of faces that coincide with axes can be disregarded. For the rest of the faces, upon lengthy but basic calculations, we also show that the solution goes inside the domain. The results are also true for *b*_0_ ≥ 3*/*4. In this case, the angle of the trapezoidal face of the domain is modified such that it is contained in the plane *Q* + *A − aN*, with constant *a<* 1. For *b*_0_ *>* 1*/*2, the positive steady state is stable: all three eigenvalues of the linearised system are real and negative. We find that the characteristic polynomial corresponding to the linearisation matrix of the reduced system (1_*\**_) has all coefficients negative. Thus, by the Routh-Hurwitz stability criterion we obtain that all real parts of the matrix eigenvalues are negative, and thus the steady state is stable.

### Parameter estimation

In order to see how our models are able to describe the dynamics observed in data, we performed parameter estimation. The proliferation rates *p*_*A*_ = 0.95 and *p*_*T*_ = 0.81 were kept constant, assuming a cell cycle length of 17h for aNSCs and 20h for TAPs [12]. In a first step, the parameters to be estimated comprised of *r*_0_, *K, b*_0_, *β, δ,N SC*_0_, where *NSC*_0_ represents the total number of stem cells (*Q* + *A*) at time *t* = 0. Having performed a large number of estimations the values for the last two parameters were similar most of the times, also among different scenarios. Therefore, for simplicity, we fixed *δ* = 0.19 and *NSC*_0_ = 1900. As a result, the parameters estimated comprise of *r*_0_, *K, b*_0_ and *β* for WT data. In the case of the IFNAGR KO experiments, the parameters *r*_1_ and *p*_*T*_ were additionally estimated. For the young mice treated with TMZ, *p*_*T*_ was fitted, together with the death rate *d* and scaling factor *ρ* (see section “Model of TMZ treatment” of Methods), while keeping the rest fixed. For old mice, both parameters *r*_0_ and *p*_*T*_, as well as *d* and *ρ* were fitted.

The optimization was based on weighted least squares. A cost function was defined as the sum of squared differences between the values *Q* +*A, A/*(*Q* +*A*), Σ*i T*_*i*_ and *N* given by the ODE system at the *j* ∈ {1, …, *n*_*t*_} time points at which data is available and the corresponding data values for each mouse (*n*_*j*_ total data points at time point *j*), divided by the variance among mice at the same time point:

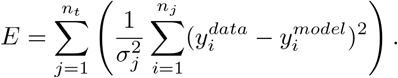

The cost function was then minimized by using the `fmincon` routine in MathWorks MATLAB software, with the sequential quadratic programming (`‘sqp’`) algorithm. The algorithm was run multiple times starting from 500 initial guesses for each model hypothesis, and the parameter set with the smallest cost function value out of the 500 multi-start points was selected.

Furthermore, after performing parameter estimation using the constrained Gauss-Newton method, a sensitivity analysis is conducted to evaluate the uncertainty of the estimated parameters. Employing the generalized inverse of the Jacobian matrix of the underlying parameter estimation problem allows for the construction of a Taylor expansion to describe how the parameter estimates respond to perturbations in the measurement errors. This expansion enables the derivation of linearized (ellipsoidal) or quadratic confidence regions and a linear or quadratic estimate of the covariance matrix of the parameters [47]. The square roots of the diagonal elements of the covariance matrix represent the standard deviations of the estimated parameters, and the off-diagonal elements indicate the correlations between the parameters. These confidence regions are local approximations of the nonlinear confidence regions defined by the maximum likelihood ratio [48] and may underestimate uncertainty in regions where the model exhibits strong nonlinearity. Nevertheless, they provide valuable insight into parameter uncertainty, which can be refined using the Bayesian approach, and can be computed efficiently.

#### Bayesian approach to sensitivity and uncertainty quantification

To quantify the sensitivity of our model’s dynamics to variations in initial conditions and parameter values, we employed Bayesian methods. Specifically, this approach was applied to three model scenarios discussed in the paper Scenario *r*(*Q, A*), *bA*(*Q*) (Eq. (10)) Scenario *r*(*Q* + *A, Q* + *A*), *b*(*Q* + *A*) (Eq. (12)) and the Delta-Notch Scenario *r*(*Q, Q* + *A*), *b*(*Q*) (Eq. (16)).

Due to the limited experimental data, which prohibited extracting reliable information about the statistics of the measurements *y*_*i*_ we assumed that each measurement *y*_*i*_ at a given time point follows a Gaussian distribution. Moreover, because these measurements were obtained from different mice, we considered measurements at different time points to be statistically independent. Model simulations revealed a rapid temporal decrease in the variance of trajectories generated from varying initial conditions. Accordingly, we assumed an exponential decay in variance of the form

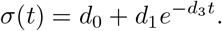

This approximation allowed us to interpolate the variance for most sparsely sampled measurements *y*_*i*_ and provided a rough estimation of the distributions for the four measured quantities in the initial conditions at *t* = 0 *Q* + *A*, 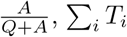, and *N*. We used these assumptions to quantify the variability of model solutions with respect to initial conditions. In our numerical simulations, these four quantities were sampled from their respective Gaussian distributions, and the components of the initial state were adjusted to satisfy the sampled proportions.

Under the assumption of normally distributed data, we expressed the discrepancy between model and experimental data *y*_*i*_ as a negative log-likelihood function:

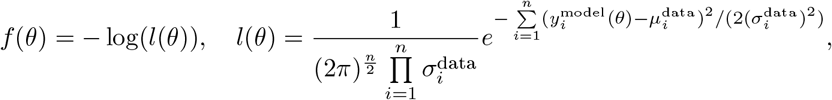

where *θ* denotes the vector of model control parameters, and 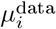 and 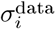 are the mean values and variances estimated from the experimental data using the method described above. The values 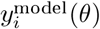 are generated from model trajectories corresponding to realisations of randomised initial conditions. Although the random sampling of initial conditions makes *f*(*θ*) inherently stochastic, numerical evaluations across different parameter values revealed that the variability for a fixed parameter vector *θ* is minimal (in general, less than 1%), as shown in Fig. 7E-E’. Therefore, we evaluated *f* (*θ*) by simulating the model multiple times with different realizations of initial conditions and next averaging the results.

Subsequently, we minimised *f* (*θ*) using the previously obtained point estimates as initial guesses (values given in Supplementary Table 1). In all cases, the optimized parameter values closely aligned with those derived from the LSQ cost function.

For uncertainty quantification, we employed the Adaptive Metropolis algorithm [41]. Here, the Markov Chain Monte Carlo (MCMC) sampler was initialised with the optimised parameter estimates. The resulting posterior distributions revealed that the parameters *r*_0_, *r*_1_, and *K* exhibit very large variability, indicating significant uncertainty in their estimates, whereas the posterior distributions for *b*_0_ and *β* were comparatively more concentrated, reflecting higher identifiability under the experimental conditions.

### Model selection

Model selection is performed by assigning each model a score that combines its goodness of fit with a penalty on the number of parameters to avoid overfitting. The Akaike Information Criterion, with its formulation for weighted least squares, and its correction for small samples, is one of the common scores used, defined as below [49], respectively.

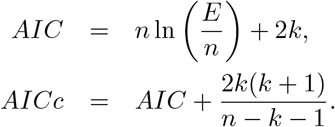

Here, *k* is the number of fitted parameters, *n* is the size of the data, and *E* represents the value of the minimized objective function. A smaller *AICc* value represents a better model, in the sense of both how well it fits the data and how parsimonious it is. In model selection, the actual *AICc* value of a model carries no real significance unless compared to those of other models. In other words, the important values to inspect are

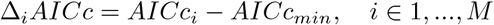

representing the difference between each of the *M* models and the best-valued one. Despite only one model having Δ_*i*_*AICc* = 0, other models with very small values should also not be discarded. As there is no accurate way of deciding where the boundary between (equally-) acceptable models and bad ones should be drawn, one can resort to computing an additional metric, namely the Akaike weights. These weights can quantify how likely a model is to be the correct one, and are defined as

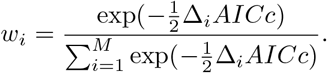

All scenarios considered in this paper have the same number of parameters and therefore we might simply compare the *E*-values of the minimized cost functions. However, for consistency and possible future generalization we compare the Δ_*i*_*AICc* and Akaike weights *w*_*i*_. The values of these scores for each hypothesis considered can be found in the Supplement.

### Model of TMZ treatment

Ta ing into account that the data from TMZ-perturbed mice consider the numbers of proliferating cells from sections of the ventricular-subventricular zone, as opposed to cell counts from the whole region, the TMZ data need to be scaled up to compare to the WT data. This is done by fitting an additional parameter *ρ* that represents the scaling factor of the section-data with respect to the whole region, based on the so-called saline data recorded few days before the TMZ experiments (56 and 656 days of age in young and old mice, respectively). The scaling factor *ρ* thus brings the values of saline data up to the order of magnitude of WT data.

Additionally, since the death of proliferating cells is not instant upon TMZ treatment, we model an exponential decay of cell counts that considers a constant death rate *d* of aNSCs (*A*) and TAPs (*T*_*i*_, *n* = 3) due to chemotherapy during the short time-span of administering the drug. The resulting ODE system is shown in Eq. (17).

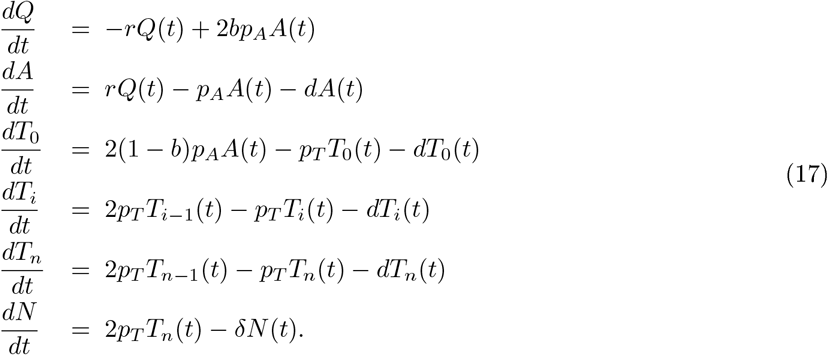

The death parameter *d* is also estimated, when fitting the model to the experimental data. Specifically, for the period before treatment the healthy system (1) in Fig. 1B is simulated, during the treatment the model (17) is used, and after the treatment window the WT model is again used. The combined solution from these three regimes is then compared to our data points and the parameters are fitted.

### Data and code availability

All original code has been deposited at Zenodo under the DOI 10.5281/zenodo.14944969 and is publicly available as of the date of publication. Any additional information required is available from the corresponding authors upon request. The data sets analyzed during the current study are either publicly available or can be obtained from the corresponding author of the original study that generated the data set [12, 36].

## ACKNOWLEDGMENTS

The project was supported by the European Research Council (ERC) under the European Union’s Horizon 2020 research and innovation programme (synergy project PEPS, no. 101071786), and by the Deutsche Forschungsgemeinschaft (DFG) within the Collaborative Research Centre SFB1324 (B05 of AMC and B06 of AM). The authors than Prof. Mare Kimmel for his feedbac and invaluable advice on manuscript preparation.

## AUTHOR CONTRIBUTIONS

D-PD: Model development, Algorithm implementation, Interpretation of the modelling results, Manuscript preparation - original draft; FZK: Mathematical analysis, Manuscript preparation - review and editing; AK: Bayesian inference, Sensitivity and uncertainty quantification, Manuscript preparation - review; LF: Multiple-shooting optimization, Optimal experimental design; EK: Supervision, Multiple-shooting optimization, Optimal experimental design, Manuscript preparation - review; AM- : Conceptualization, Supervision, Interpretation of the modelling results, Manuscript preparation - review; AM-C: Conceptualization, Supervision, Model development, Interpretation of the modelling results, Manuscript preparation - review and editing;

