## Supplemental Information for "Unraveling Regulatory Feedback Mechanisms in Adult Neurogenesis Through Mathematical Modelling"

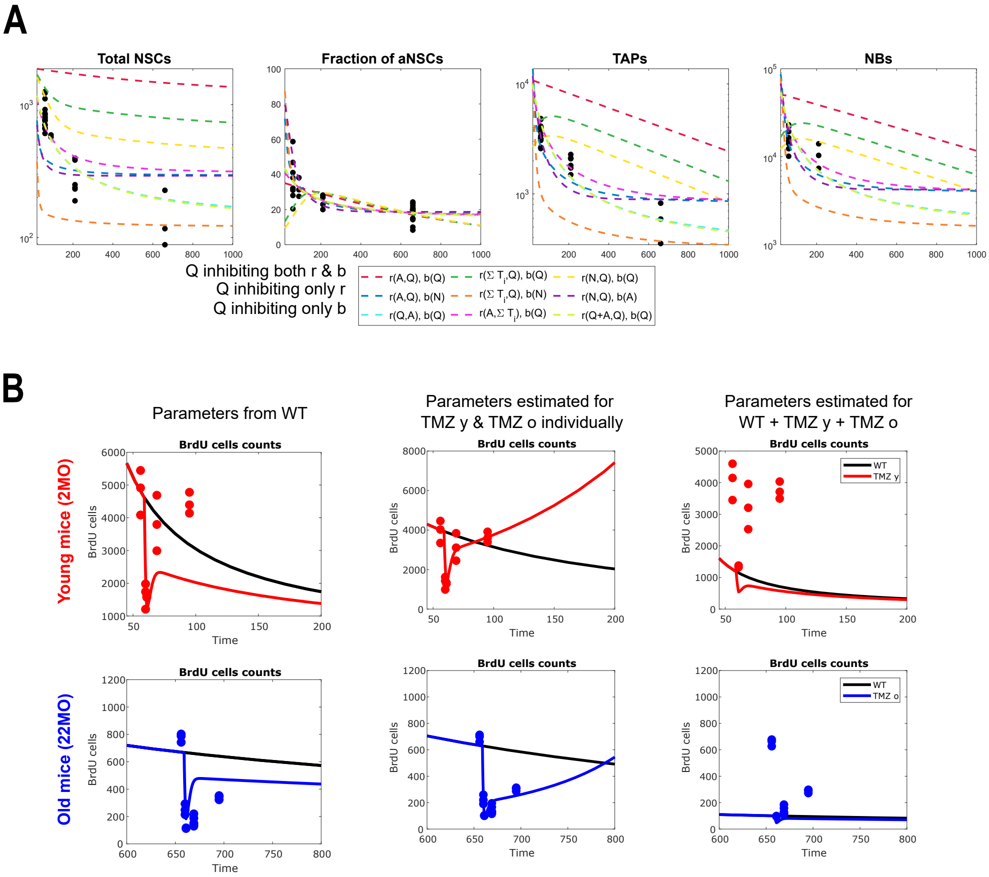


**Supplementary Figure 1:** Examples of bad fits to WT and TMZ data: **A.** Scenarios in which either Q inhibits both $r$ and $b$, only $r$ or only $b$. The estimated $b_0 > 1/2$ for all these, so the solutions converge to the positive steady state. **B.** Results of *in silico* TMZ treatment with parameters estimated for WT (left); all parameters estimated individually for young and old TMZ data (middle); and parameters estimated for fitting all data together (from WT, TMZ young and TMZ old mice). Red represents TMZ-treated young mice, blue corresponds to TMZ-treated old mice and black to WT mice without treatment. The scenario shown is that with system parameters $r(Q, A)$ and $b(A)$ (3), but the same behaviour is seen for all of the five best-scoring hypotheses (3)-(7).

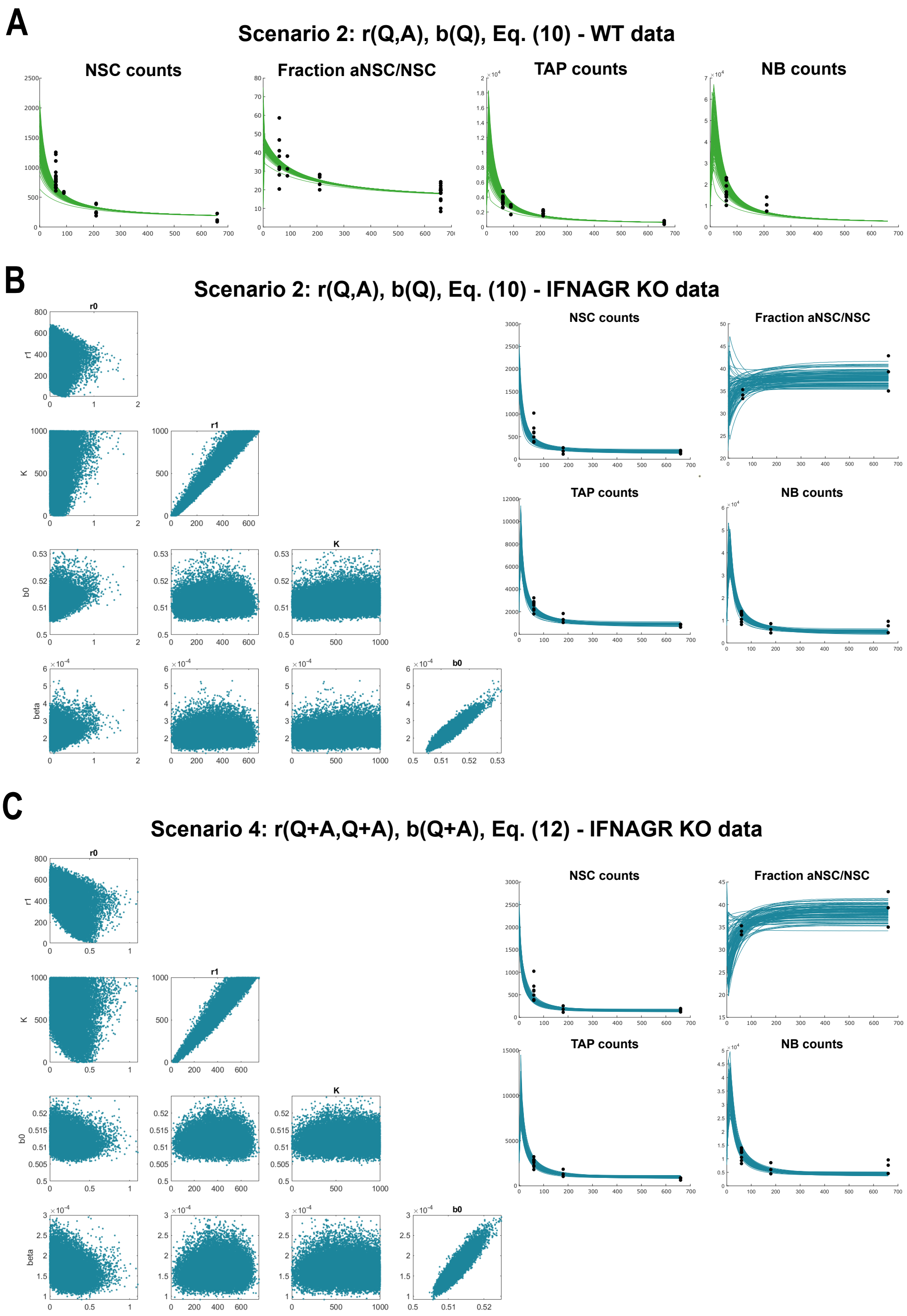


**Supplementary Figure 2:** Uncertainty and sensitivity quantification: **A.** Model simulations for Scenario 2 (Eq. (4)), with parameters estimated for WT data using the least-squares method, shown in Table S1, starting from a Gaussian distribution of initial conditions with mean and variance interpolated from the data (see Methods for details). All simulations quickly converge to the same dynamics, regardless of the starting values. **B.** Posterior distributions and correlations for model parameters, for Scenario 2, $r(Q, A)$, $b(Q)$, given by Eq. (10), applied to data from IFNAGR KO mice. Right-hand side inset shows the model trajectories starting from numerous initial conditions and with parameter estimates from the MCMC results (see Methods for details). **C.** Similar posterior distribution and treajectories plots as in **B.**, for Scenario 4: $r(Q + A, Q + A)$, $b(Q + A)$, given in Eq. (12).

**Supplementary Table 1**: Parameter values for the six scenarios

| Scenario | r0_wt | K_wt | b0_wt | beta_wt | r0_ifnko | r1_ifnko | K_ifnko | b0_ifnko | beta_ifnko | pT_ifnko | pT_TMZ_y | d_TMZ_y | r0_TMZ_o | pT_TMZ_o | d_TMZ_o | rho_TMZ |
|---|---|---|---|---|---|---|---|---|---|---|---|---|---|---|---|---|
| r(Q,A), b(A) | 2.3583 | 1774.3 | 0.49994 | 0.00011306 | 221.03 | 0 | 301.94 | 0.51611 | 0.0005284 | 1.052 | 0.44291 | 0.66398 | 0.22357 | 0.18146 | 0.96003 | 39.128 |
| r(Q,A), b(Q) | 2.4376 | 1808.5 | 0.50557 | 9.0097E-05 | 191.17 | 0.0030268 | 253.77 | 0.51106 | 0.0002341 | 1.0313 | 0.44968 | 0.71346 | 0.22986 | 0.18068 | 0.95958 | 38.744 |
| r(Q,A), b(N) | 2.2707 | 1735.5 | 0.49952 | 1.195E-06 | 191.91 | 0.019638 | 256.84 | 0.51306 | 5.4708E-06 | 1.1016 | 0.45263 | 0.67971 | 0.21505 | 0.18101 | 0.95967 | 39.985 |
| r(Q+A), b(Q+A) | 1.5015 | 1252.3 | 0.50355 | 5.0994E-05 | 315.52 | 0.21125 | 415.88 | 0.51317 | 0.00016873 | 1.0353 | 0.43631 | 0.66279 | 0.16984 | 0.18093 | 0.96058 | 40.516 |
| r(Q,T0), b(N) | 2.4994 | 1857.9 | 0.49952 | 1.2342E-06 | 227.09 | 0 | 316.7 | 0.51284 | 5.4203E-06 | 1.0502 | 0.44646 | 0.68597 | 0.23758 | 0.18032 | 0.95958 | 38.926 |
| r(Q,Q+A), b(Q) | 5.5205 | 3966.1 | 0.50542 | 8.8927e-05 | 521.13 | 0.077494 | 712.5 | 0.51192 | 0.00025174 | 1.042 | 0.45268 | 0.71795 | 0.52731 | 0.18069 | 0.95946 | 38.63 |

**Supplementary Table 2**: Akaike scores for each scenario

| Scenario | AICc | ∆AICc | Akaike weights |
|---|---|---|---|
| **r(Q,A), b(A)** | -1427.537237 | 0.749237896 | 23.0871787 |
| **r(Q,A), b(Q)** | -1428.286475 | 0 | 33.57884908 |
| **r(Q,A), b(N)** | -1424.658985 | 3.627490398 | 5.474774993 |
| **r(Q+A), b(Q+A)** | -1428.148514 | 0.137960875 | 31.34064882 |
| **r(Q,T0), b(N)** | -1425.007986 | 3.278489326 | 6.518548405 |

**Supplementary Table 3**: Akaike scores for the Delta-Notch and two “best” scenarios for WT data

| Scenario | AICc | ∆AICc | Akaike weights |
|---|---|---|---|
| **r(Q,A), b(Q)** | -728.49 | 0.016838 | 34.036 % |
| **r(Q+A), b(Q+A)** | -728.34 | 0.16275 | 31.641 % |
| **r(Q,Q+A), b(Q)** | -728.51 | 0 | 34.323 % |
